# Murine uterine gland branching is necessary for gland function in implantation

**DOI:** 10.1101/2023.11.01.565233

**Authors:** Katrina Granger, Sarah Fitch, May Shen, Jarrett Lloyd, Aishwarya Bhurke, Jonathan Hancock, Xiaoqin Ye, Ripla Arora

**Affiliations:** Department of Obstetrics, Gynecology and Reproductive Biology, Michigan State University; Institute for Quantitative Health Science and Engineering, Michigan State University; Department of Physiology and Pharmacology, College of Veterinary Medicine, University of Georgia; Interdisciplinary Toxicology Program, University of Georgia

**Author notes:** Equal contribution (Katrina Granger, Sarah Fitch). **Corresponding Author** Ripla Arora Associate Professor, Department of Obstetrics, Gynecology and Reproductive Biology Institute for Quantitative Health Science and Engineering Michigan State University.

## Abstract

Uterine glands are branched, tubular structures whose secretions are essential for pregnancy success. It is known that pre-implantation glandular expression of leukemia inhibitory factor (LIF) is crucial for embryo implantation, however contribution of uterine gland structure to gland secretions such as LIF is not known. Here we use mice deficient in estrogen receptor 1 (ESR1) signaling to uncover the role of ESR1 signaling in gland branching and the role of a branched structure in LIF secretion and embryo implantation. We observed that deletion of ESR1 in neonatal uterine epithelium, stroma and muscle using the progesterone receptor *Pgr^Cre^* causes a block in uterine gland development at the gland bud stage. Embryonic epithelial deletion of ESR1 using a mullerian duct Cre line - *Pax2^Cre^*, displays gland bud elongation but a failure in gland branching. Surprisingly, adult uterine epithelial deletion of ESR1 using the lactoferrin-Cre (*Ltf^Cre^*) displays normally branched uterine glands. Intriguingly, unbranched glands from *Pax2^Cre^ Esr1^flox/flox^* uteri fail to express glandular pre-implantation *Lif,* preventing implantation chamber formation and embryo alignment along the uterine mesometrial-antimesometrial axis. In contrast, branched glands from *Ltf^Cre^ Esr1^flox/flox^* uteri display reduced expression of glandular *Lif* resulting in delayed implantation chamber formation and embryo-uterine axes alignment but deliver a normal number of pups. Finally, pre-pubertal unbranched glands in control mice express *Lif* in the luminal epithelium but fail to express *Lif* in the glandular epithelium even in the presence of estrogen. These data strongly suggest that branched glands are necessary for pre-implantation glandular *Lif* expression for implantation success. Our study is the first to identify a relationship between the branched structure and secretory function of uterine glands and provides a framework for understanding how uterine gland structure-function contributes to pregnancy success.

## INTRODUCTION

Uterine glands are key to embryo implantation and pregnancy success. In viviparous mammals in the absence of yolk, uterine glands support the pregnancy and embryo development until the placenta forms (1). Uterine glands are present in humans, rodents, sheep, pigs, and horses among other studied mammals (2). Uteri of neonatal mice are devoid of glands at birth. At postnatal day (P) 5, gland buds protrude off the uterine lumen on the anti-mesometrial (AM) and lateral sides of the murine uterus and extend into the surrounding stroma Glands are branched in the pubertal non-pregnant and pregnancy stages^1^. Uterine glands are categorized as exocrine glands, however, unlike other branched exocrine glands such as salivary and mammary glands, mechanisms of uterine gland branching have not been identified Uterine glands facilitate the secretions of vital factors including the key cytokine Leukemia inhibitory Factor (LIF), that is essential for embryo implantation (5, 6). While it is known that exocrine gland structure contributes to function in the case of meibomian glands, mammary glands and salivary glands (4), whether uterine gland structure contributes to gland secretory function is not known.

Estrogen (E2) signaling facilitates the growth and development of the uterus as suggested by various estrogen receptor deletion models. Estrogen receptor has two isoforms, ESR1 and ESR2, which are encoded by separate genes located on different chromosomes. Both receptor isoforms are nuclear receptors and act as ligand-activated transcription factors that can alter target gene transcription (7). Female mice with a whole body ESR1 deletion are infertile (8). These females fail to ovulate and exhibit hypoplastic uteri that fail to display cyclic changes (9). In contrast, females with a whole body ESR2 deletion have sub-fertility with a reduced litter size (8). Mice with conditional deletion of *Esr1* in different compartments have been generated to highlight the cell-type-specific contributions of ESR1 signaling in the uterus. Deletion of ESR1 in the neonatal epithelium, stroma and muscle, using progesterone receptor driven *Pgr^Cre^*, and in embryonic epithelium using *Wnt7a^Cre^* results in infertility (10, 11). However, implantation has not been assessed in *Pgr^Cre^* model and epithelial deletion of ESR1 using *Wnt7a^Cre^* results in protease-mediated embryo death in the oviduct (12). Embryo transfer studies into pseudopregnant uteri of the latter mice suggest that embryo implantation is not supported in this model (10).

Gland development is variably responsive to ovarian E2 depending on the species. In the neonatal pig, gland development is both estrogen receptor 1 (ESR1)-dependent and sensitive to E2 levels (13, 14). Unlike the pig, in sheep, ESR1-inhibition does not impact gland initiation (15) but reduces number of glands in the intercaruncular area, and reduces branches and coils in the uterine glands (15). In neonatal mice, the initiation of gland formation is thought to be ovary-, adrenal-, and steroid-independent (16, 17). However, treatment with Genistein (an ESR1 agonist) during the neonatal period prevents gland development and results in implantation failure (18). Gland structure in *Esr1*-deficient mice has not been assessed.

Mouse mutants that display an absence of glands or key glandular secretion LIF display implantation and decidualization failure (19). Examples include mice-deficient in *Wnt7a* (20); *Foxa2* (21, 22); Progesterone-induced gland knockout (23) and *Lif* (5). The luminal epithelium, glandular epithelium, and stromal compartment all possess the ability to express *Lif*, and the coordination of LIF secretion among these compartments presumably happens in accordance with steroid hormone levels during both the mouse estrous cycle and early pregnancy (24). ESR1 signaling is critical for *Lif* induction. Ovulatory E2 coincides with ESR1 and *Lif* expression in the uterus (25, 26) and prior to implantation, uterine glands express ESR1 and LIF (26). A large dose of E2 has the effect of inducing robust *Lif* expression, and this response is absent in ESR1 knockout mice (10, 27). However, the compartment in which *Lif* is expressed (luminal or glandular epithelium) when stimulated by a large dose of E2 remains unknown. *Wnt7a^Cre^* deletion of ESR1 results in a reduction of *Lif* and failed oil-mediated artificial decidualization, and exogenous *Lif* supplementation rescues artificial decidualization in this model (11). This highlights the need for epithelial ESR1 signaling for pre-implantation *Lif* production in early pregnancy.

Studies have demonstrated that there is a link between ESR1 signaling and mammary gland branching such that both complete abrogation of ESR1 signaling and epithelial-specific deletion of ESR1 result in significantly reduced mammary gland branching and consequently diminished lactational function (28–30). While the role of ESR1 signaling has been separately studied during uterine development and for pre-implantation *Lif* expression, a connection between gland structure and function is yet to be made. In this study, we determine the relationship between estrogen signaling, development of a branched gland structure and pre-implantation glandular LIF production. We determine that neonatal stromal ESR1 signaling is key to gland elongation and pubertal ESR1 signaling is necessary for gland branching. We also uncovered that unbranched and ESR1-deficient glands fail to express *Lif*, whereas branched ESR1-depleted glands can produce enough LIF to support embryo implantation and pregnancy. Finally, we show that pre-pubertal uteri support *Lif* expression in the luminal epithelium but not in the unbranched glands, suggesting that gland structure (branching) is necessary for gland function (glandular LIF production and implantation).

## MATERIALS AND METHODS

### Animals

*Esr1^flox/flox^* mice were provided by Dr. Pierre Chambon (31). These mice were bred with *Pgr^Cre^*(32)*, Pax2^Cre^* (33), or *Ltf^Cre^* (34) mouse lines to generate tissue and time specific deletion of ESR1 (Table 1). *Esr1^flox/flox^*females were used as controls. For pregnancy studies females were mated with fertile CD1 males and the appearance of a vaginal plug was identified as gestational day (GD) 0 1200h. Adult females > 6 weeks were used for *Pgr^Cre^ Esr1^flox/flox^*(hereafter referred to as PER) and *Pax2^Cre^ Esr1^flox/flox^* (hereafter referred to as XER) deletion models. In the *Ltf^Cre^ Esr1^flox/flox^* deletion model (hereafter referred to as LER), adult females aged 10-12 weeks were used. For pups study, LER virgin females were set up with CD1 males, monitored for pregnancy, and the number of live pups were recorded at birth. *Wnt7a^Cre^ Esr1^flox/flox^* mice (hereafter referred to as WER) were used for analysis of uterine gland structure to compare with XER mice. These *Esr1^flox/flox^* mice were generated by Dr. Kenneth Korach (35). Both the *Esr1^flox/flox^* mice used in the study have exon 3 of the *Esr1* gene flanked by LoxP sites and result in complete absence of the ESR1 protein. To detect implantation sites, female mice at GD4 1200h mice were injected intravenously or retroorbitally with blue dye prior to dissection. All animals were maintained and handled according to the Michigan State University Institutional Animal Care and Use Committee guidelines.

**Table 1.** Genotypes and abbreviations used in the study.

| Mouse Line | Abbreviation | Uterine Cell-type in which deletion occurs | Time of deletion |
| --- | --- | --- | --- |
| *Pgr*<sup>Cre</sup> *ESR1*<sup>flox/flox</sup> | PER | Epithelium, Stroma, Circular Muscle | Neonatal |
| *Pax2*<sup>Cre</sup> *ESR1*<sup>flox/flox</sup> | XER | Epithelium | Embryonic |
| *Wnt7a*<sup>Cre</sup> *ESR1*<sup>flox/flox</sup> | WER | Epithelium | Embryonic |
| *Ltf*<sup>Cre</sup> *ESR1*<sup>flox/flox</sup> | LER | Epithelium | Adult |

### LIF, Hormone and Inhibitor Treatments

For *Lif* rescue experiments, XER mice were injected intraperitoneally at 1000h and 1800h on GD3 with either PBS or 10µg recombinant *Lif* (554006, BioLegend, San Diego, CA) followed by blue dye injection and dissection at GD4 1200h. Alternatively, one uterine horn of XER mice was injected intraluminally with 1µg recombinant *Lif* (BioLegend, 554006) at 1300h on GD3. The partner horn was left untreated as a control. Mice were then dissected on GD4 1200h following blue dye injectionFor exogenous hormone treatments, 17D-estradiol (E2) (E8875, Sigma-Aldrich, St. Louis, MO, USA) and progesterone (P4) (P0130, Sigma-Aldrich, St. Louis, MO, USA) were dissolved in sesame oil and two injection schemes were used. In the first method, control postnatal day (P) 21 mice were subcutaneously injected with either 100ng E2 or sesame oil at 1200h on P21 and 0900h on P22 before dissection at 1200h on P22. In the second method, control P21 mice were subcutaneously injected with 100ng E2 at 1200h on P21 and P22. On P23, mice were injected with 1mg P4 + 6.7ng E2 at 0900h and subsequently dissected at 1200h.

### Cryoembedding, Cryosectioning and immunostaining

For cryoembedding, uterine horns were dissected and fixed in 4% paraformaldehyde overnight at 4°C. The next morning, uteri were washed three times 5 min each with PBS and left upright in a solution of 10% sucrose overnight. Uteri were then transferred into 20% and 30% sucrose solutions, respectively, for 2-3 h each. Finally, uteri were embedded longitudinally in Tissue-Tek OCT (Andwin Scientific, 45831) and stored at -80°C. Tissues were cryosectioned at 7µm, mounted on glass slides (Fisher, 1255015), and stored at -20 until immunostaining. For ESR1 immunostaining, antigen retrieval was performed by washing slides in PBS before transferring to 1x citrate buffer solution (Life Sciences, 005000) and boiling in a beaker filled to ¾ in the microwave for 10 min. Following this, slides were washed three times 5 min each with PBS, blocked with 2% powdered milk in PBS + 1% Triton (PBT), and left in primary antibodies overnight at 4°C. The next day, slides were washed three times 5 min each with PBS, stained with secondary antibodies for 1 h, washed with PBS again, and sealed with a coverslip and nail polish.

### Confirmation of Pax2 Cre-lineage

Pax2 lineage was confirmed by breeding *Pax2^Cre^* mouse line with *Gt(ROSA)26Sor^tm^*^4^*^(ACTB-tdTomato,-EGFP)Luo/J^*also referred to as *ROSA^mT/mG^* (Jackson Labs, 007576) reporter mice. Uteri from P21 pups were dissected and embedded for cryosectioning. Cryosections were imaged for endogenous membrane GFP and membrane tomato signal. Some cryosections were additionally stained with primary antibody for CD31 to identify blood vessels (see tissue section immunofluorescence).

### Whole-mount and tissue section immunofluorescence

Whole-mount immunofluorescence was performed as previously described (36) Uteri were dissected from mice and fixed in DMSO:methanol (1:4). For immunostaining, uteri were rehydrated in methanol:PBT (1% Triton X-100 in PBS) (1:1) for 15 min, washed in PBT for 15 min and incubated in blocking solution (2% powdered milk in PBT) for 1 h at room temperature. Uteri were incubated with 1:500 concentration of primary antibodies diluted in blocking solution for seven to nine nights at 4°C. They were then washed twice for 15 min each with 1% PBT followed by three washes for 45 min each at room temperature. Uteri were then incubated with secondary antibodies at 4°C for two or three nights, followed by one 15 min and three 45 min washes with 1% PBT and dehydration in 100% methanol for 30 min. Uteri were then bleached overnight at 4°C in a solution of 3% H_2_O_2_ in methanol. Finally, the samples were washed in 100% methanol for 1.5 h and cleared in BABB (1:2, benzyl alcohol:benzyl benzoate) (Sigma-Aldrich, 108006, B6630).

### Antibodies

Primary antibodies used include: mouse anti-ESR1 (Thermofisher, MA5-13191; Abcam, ab93021; 1:200,), rat anti-CDH1 (M108, Takara Biosciences, 1:500), rabbit anti-FOXA2 (Abcam, ab108422; 1:500, 1:200), PTGS2 (Abcam, ab16701; 1:500), Rat anti-CD31 (BD Biosciences, B553370) and Biotin-conjugated isolectin B4 (Vector Laboratories, B1205.5, 1:200). Alexa Fluor-conjugated secondary antibodies: donkey anti-mouse 555 (1:500), goat anti-rat 647 (Invitrogen, A21247; 1:500), goat anti-rat 633 (Invitrogen, A21094; 1:500), and donkey anti-rabbit 555 (Invitrogen, A31572; 1:500), were obtained from Invitrogen, and Hoechst (Sigma Aldrich, B2261) was used to stain the nucleus. Streptavidin Dylight 633 (Invitrogen, 21844; 1:500) was used as a secondary antibody for isolectin B4.

### In situ hybridization

In situ hybridization on uterine sections was performed using the RNAscope 2.5 HD Assay-RED kit (ACD Bio, 322350), which incorporates immunofluorescence capabilities. Mm-*Lif* probe (ACD Bio, 475841) was used to detect *Lif* mRNA and immunostaining for FOXA2 was included to label uterine glands. The technique was carried out according to the protocols outlined by ACD Bio (322360-USM, MK 51-149 TN). Primary and secondary antibodies used include rabbit anti-FOXA2 (Abcam, ab108422; 1:200) and Alexa-Fluorophore donkey anti-rabbit 647 (1:500), respectively. Hoechst (Sigma Aldrich, B2261) was used to stain the nucleus.

### Confocal microscopy

Samples with whole tissue immunofluorescence, section immunofluorescence and in situ hybridization were all imaged using a Leica TCS SP8 X Confocal Laser Scanning Microscope System with white-light laser, 10× air objective (used for whole tissue) and 20× water objective (used for sections). The entire length and thickness of the uterine horn was imaged using the tile scan function with z stacks 7μm apart. For sections, z stacks 1.5μm apart were used. Images were merged using Leica software LASX version 3.5.5.

### 3D reconstruction and image analysis

Image analysis was performed using commercial software Imaris v9.2.1 (Bitplane). The confocal image (.LIF) files were imported into the Surpass mode of Imaris.

<u>For gland visualization</u>, the Surface function of Imaris was used to reconstruct 3D gland surfaces based on the FOXA2 fluorescent signal. The Imaris Vantage function was used to isolate individual glands into a comprehensive gallery for visualization.

<u>For quantitative analysis of gland length and branch numbers,</u> images of uterine horns were imported into Imaris v9.2.1 (Bitplane) with Matlab (XT) module. The surface function in Imaris was used to create 3D renderings of uterine glands from the fluorescent staining by background subtraction with the diameter of the largest sphere set to 30. With files of uterine horns stained only with Cytokeratin 8, glands were isolated manually from the surface by using the scissors tool to cut them away from the lumen. For gland length the Bounding Box OOC function in Imaris was used that determines the shortest straight-line distance from the point where the gland is connected to the uterine lumen to the furthest tip. Masks of the gland Surfaces were made to get a channel with uniform gland signal throughout. These channels were then imported to FIJI (ImageJ). Thresholding was used to binarize the isolated image and the "Fill Holes" function was used to fill in any gaps to ensure a more accurate skeleton. This 3D image was then saved as a TIFF file and imported into MATLAB. The "imread" function was used so MATLAB could read the data and a new variable was defined using the command "uint8(Skeleton3D(imbinarize(stack)))*200;". This command defines the file as a binary image, calculates the 3D skeleton of a binary volume using a thinning algorithm and increases the intensity of the signal throughout. The "saveastiff" function was then used to save this revised file as a TIFF file, which was then added to the original Imaris file as a new channel. Filaments were created manually using that channel. The number of dendrite branch points were exported from Imaris in a Microsoft Excel file for statistical analysis.

<u>Determination of embryo axis orientation</u> was carried out by identifying embryos via Hoechst signal and using the Measurements Points module in Imaris. Embryo axis can be determined by identifying the inner cell mass (ICM) or the embryonic pole and the mural trophoectoderm (abembryonic pole). Proper embryo alignment is characterized by the embryonic pole facing the mesometrial side of the uterus and the abembryonic pole facing the anti-mesometrial side of the uterus. The embryo at an implantation site on GD4 was visualized using an optical XY Orthogonal Slicer or Oblique Slicer. An XZ Orthogonal Slicer was used to define the mesometrial-anti-mesometrial (M-AM) axis and was placed at the abembryonic pole of the embryo. Using the Measurement Points module, the first point was placed on the ICM on the M-AM plane. The second point was placed on the intersection of the M-AM plane and the abembryonic pole on the XY plane. The third point was placed on the intersection of the M-AM and XY planes. The value of the angle was obtained using the Statistics function of Imaris.

For <u>quantifying *Lif* signal</u> from in situ hybridization on cryosections, the Imaris Surface function was used to construct volumetric surfaces of gland nuclei, and *Lif* signal. For quantitation of *Lif* volume per gland volume, volumetric glandular surface of the z-stack of a single cross section of uterus was constructed according to FOXA2 signal. Then, the Imaris masking function was used to create a separate channel of *Lif* signal underneath the previously constructed gland surface volume. Based on this *Lif* signal channel, volumetric *Lif* surface was created. The Statistics function of Imaris was utilized to determine gland and *Lif* surface volumes, and Microsoft Excel was implemented to calculate *Lif* volume per gland volume

### Statistical analysis

To compare numbers of implantation sites, gland length measurements of P21 treatment mice, embryo axis orientation, and *Lif* volume per gland volume, Kruskal-Wallis test with Dunn’s multiple comparisons was conducted. For analysis of the number of implantation sites/pups, gland length measurements of GD4 1200h and P21 mice, and *Lif* volume per lumen volume, Mann-Whitney test was applied. To compare if proportions were comparable for ESR1 staining and number of glands with various branches the two-proportion Z-test was used. Statistical analyses were performed using GraphPad Prism (Dotmatics) with advanced statistical analyses conducted using R Statistical Software (37). p-value <0.05 was considered significant indicating differences between comparisons.

## RESULTS

### Generation of tissue-specific ESR1 deletion mice

To assess the role of ESR1 in uterine gland structure, we generated mice with uterine specific deletion of ESR1 in a time and compartment-specific manner (Table 1). *ESR1^flox/flox^* mice bred with a *Pgr^Cre^*mouse line (*Pgr^Cre^ ESR1^flox/flox^*, hereafter referred to as PER) were generated to determine the role of neonatal epithelial, stromal, and muscle-derived ESR1. To determine the role of epithelial ESR1 in gland structure, we used a novel *Pax2^Cre^* to generate mice where ESR1 was deleted in the embryonic Mullerian duct epithelium (*Pax2^Cre^ ESR1^flox/flox^*, hereafter referred to as XER). To determine the role of epithelial ESR1 in gland structure in the post-pubertal mouse, *Ltf^Cre^* was used (*Ltf^Cre^ ESR1^flox/flox^* hereafter referred to as LER).

Since this is the first application for *Pax2^Cre^* in the uterine epithelium, we discerned the lineage of Cre expressing cells using the *ROSA^mT/mG^* (38) reporter line. We observed that Pax2 lineage labeled cells contribute to both the luminal and glandular epithelium of the uterus but are absent from the muscle and stroma on postnatal day (P) 21 (Fig. 1A). We also observed that the oviduct displays patchy expression of the lineage label (Supp. Fig. 1). In addition to the luminal and glandular epithelium, the CD31+ vascular compartment also expresses the lineage label (GFP) (Fig. 1A). Since we are perturbing ESR1 signaling using the *Pax2^Cre^,* we determined ESR1 expression in the endothelial cells at different stages of development and at pregnancy gestational day (GD) 3 1200h. Immunostaining with ESR1 and vascular marker isolectin suggested no expression of ESR1 in endothelial cells at P21, P28, proestrus stage and GD3 1200h (Supp. 2).

**Figure 1.**
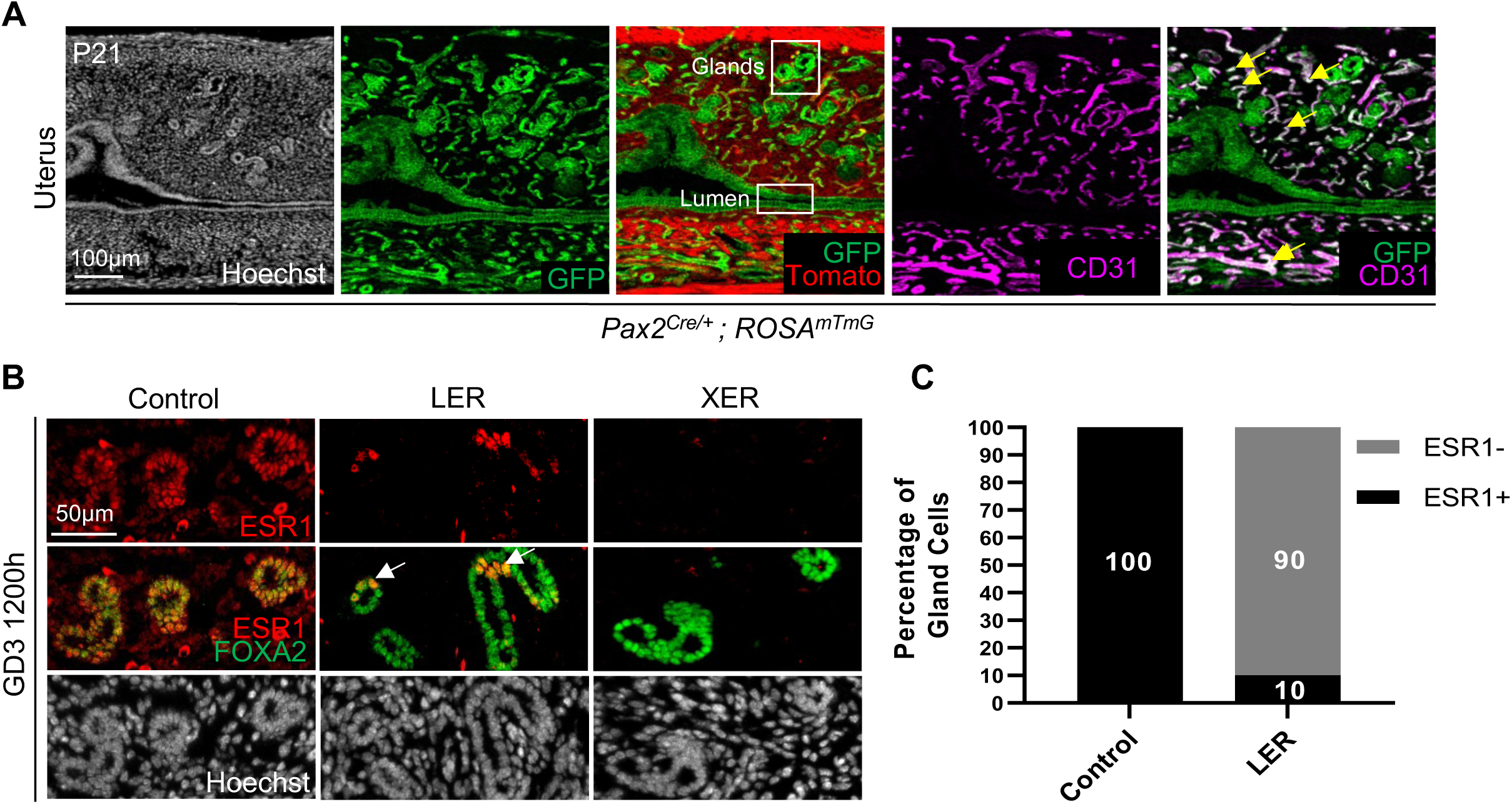
Compartment-specific excision of ESR1. (A) *Pax2^Cre/+^ ; ROSA^mTmG^* mouse uterine section at P21 with Hoechst (nuclei, white), GFP (green), Tomato (red), and CD31 (endothelial cell marker, magenta) identifying Pax2 lineage cells (green) in the luminal epithelium, glandular epithelium, and vasculature. Scale bar, 100μm; Yellow arrows indicate colocalization of GFP and CD31. (B) Uterine sections from control, LER, and XER mice identifying ESR1 (red) and FOXA2 (gland marker, green). XER mice show complete ablation of ESR1 while LER mice have few gland cells that maintain ESR1 expression (white arrowheads). Scale bar, 50μm. n=3-5 mice per genotype. (C) Quantification of percentage of ESR1+ gland cells in LER glandular epithelium compared to controls. Controls (n=4 mice, 9 sections, 2848 gland cells); LERs (n=5 mice, 17 sections, 3250 gland cells). Two-proportion Z-test determined that the differences in proportions of gland cells expressing ESR1 between control and LERs was statistically significant.

To confirm ESR1 deletion in the glandular epithelium of LER and XER mice, immunological staining with ESR1 and glandular marker FOXA2 was performed (Fig. 1B). While XER mice displayed no expression of ESR1, 10% of LER glandular epithelial cells maintained ESR1 expression at GD3 1200h (Fig. 1C), suggesting the *Ltf^Cre^* deletion of ESR1 is not a 100% complete.

### Embryonic epithelial ESR1 deletion compromises gland branching

Uterine glands in mice develop in a sequential manner from bud to elongated phase in the pre- pubertal period, eventually becoming branched in adulthood. We sought to analyze gland structure in ESR1 deletion mice at different stages. At P21, a stage when majority of the glands are unbranched (3), PER and XER mice display gland buds that are comparable to controls

(Fig. 2A, Supp. Fig. 3). When we assessed the structure of uterine glands at diestrus, a non-pregnant but pubertal stage, as observed previously glands were branched in the control uteri (36). While glands of LER appeared similarly branched to controls, glands of XER mice displayed unbranched glands (Fig. 2B). To assess gland structure during pregnancy, we analyzed GD 4 1200h uterine glands (Fig. 2C). Similar to the diestrus stage, we observed that control and LER glands were branched whereas XER glands were long but unbranched and PER glands appeared as gland buds. Using an image segmentation algorithm, we quantified gland branching to support our qualitative analysis. At P21 8% of control glands display 1-3 branches whereas neither control nor PER or XER glands display any glands with >3 branches. (Fig. 2D). At GD4 1200h, we observed an age dependent effect on gland branching. For controls at 7 weeks, 9% of glands displayed >3 branches and this number increased to 40% when controls aged >11 weeks were analyzed. Similarly, LER mice that are only analyzed at >11 weeks (to allow sufficient CRE mediated ESR1 excision), 45% of glands display >3 branches. Glands in XER and PER mice failed to display >3 branches irrespective of age evaluated (Fig. 2E, age range 7-14 weeks). In addition to gland branching, we also observed that while gland length was similar in controls, PERs, and XERs at P21, gland length was greatly reduced in GD4 1200h PERs and XERs (Supp. Fig. 4). However, LERs displayed gland length similar to controls at GD4 1200h (Supp. Fig. 4).

**Figure 2.**
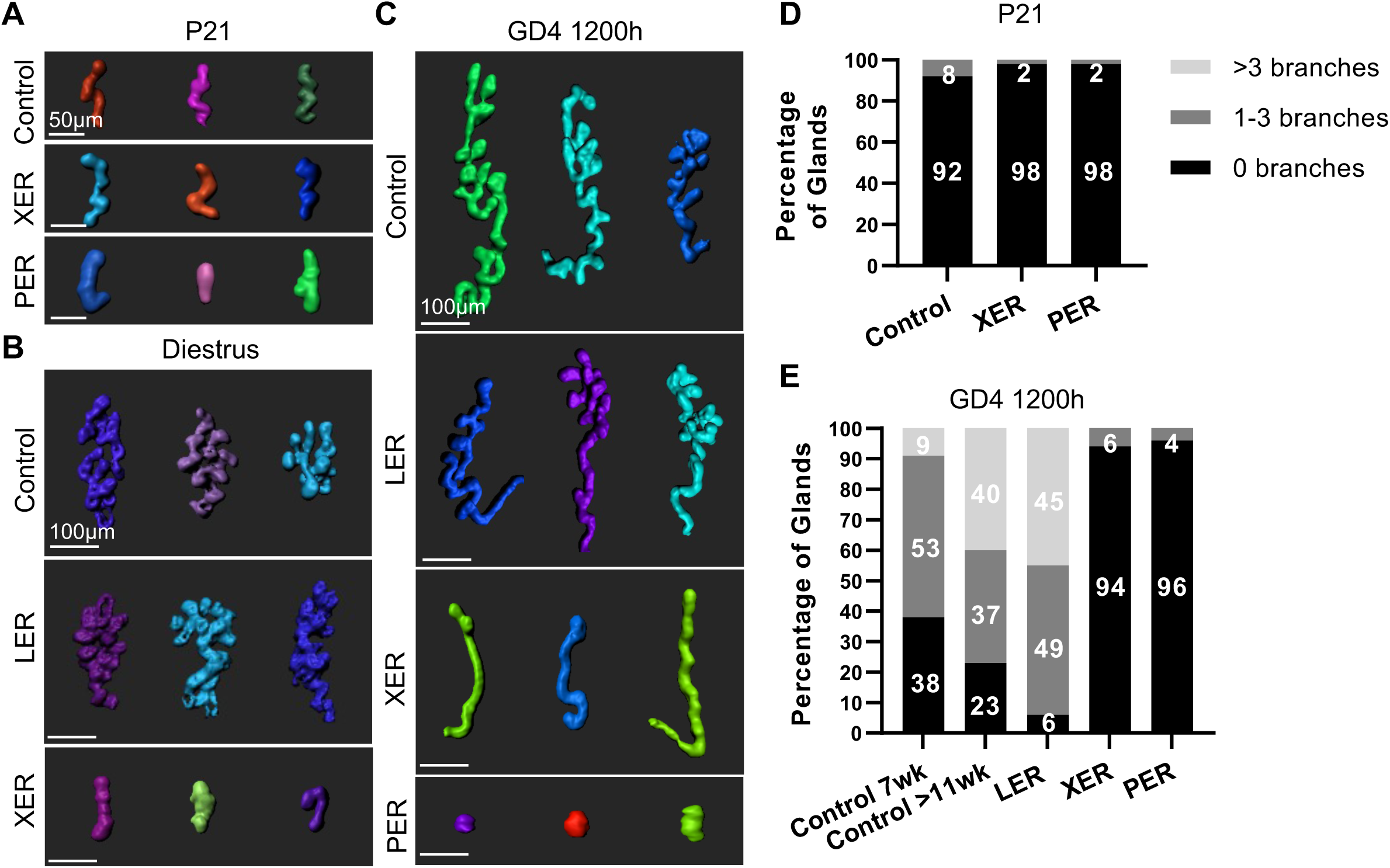
Structural phenotypes of ESR1 deletion glands during development and pregnancy. (A-C) Representative 3D reconstructions of glands at P21 (A), diestrus (B), and GD4 1200h (C). Scale bar: 50μm in A and 100μm in B and C. n=3 mice per genotype. At P21, XER and PER glands show comparable branching patterns to controls, but at GD4 1200h XER and PER glands are unbranched while control and LER glands are branched. (D-E) Quantitative analysis of percentage of glands with 0, 1-3, and >3 branches at P21 (D) and GD4 1200h (E). n=3 mice per genotype. 75-400 glands analyzed per mouse. Two-proportion Z-test was used to determine statistical differences between groups for percentage of branched glands. At GD4 1200h, gland branches between control and XER glands, and control and PER glands are statistically significantly different but there is no significant difference between control and LER or between XER and PER glands.

To confirm if gland branching defects in XERs were due to ESR1 deletion in embryonic epithelium or the vasculature, we employed another Mullerian duct Cre *Wnt7a^Cre^* to delete ESR1 in the embryonic epithelium. Similar to previous reports, we observed FOXA2 expression in the luminal epithelium of *Wnt7a^Cre^ ESR1^flox/flox^* mice (hereafter referred to as WERs), at GD3 1200h (Supp. Fig. 5A) and no embryos were observed in the uterine lumen. Similar to XERs, WERs displayed highly reduced gland branching (Supp. Fig. 5B). 61% glands in WERs displayed no branching, 34% glands displayed 1-3 branches and a small proportion displayed >3 branches (5% glands) (Supp. Fig. 5C). These results suggest that while both WNT7A and PAX2 are expressed in the embryonic Mullerian duct epithelium (39), there are differences in *Wnt7a* and *Pax2* promoter driven CRE expression that results in differential excision of *Esr1* and resulting uterine gland branching. Despite this, the extent of branching in WER uterine glands was greatly reduced compared to controls. Altogether, these observations suggest that ESR1 plays an important role in determining uterine gland elongation and branching during the initial development phase.

### Branchless glands fail to support pregnancy

To determine if gland branching defects result in functional deficits, we evaluated ESR1-deficient mice at GD3 1200h, GD4 1200h, GD4 1800h and GD5 1200h. Since WERs show a failure of embryo development in the oviduct, we wanted to first confirm that embryos did indeed enter the uterine lumen in XERs. At GD3 1200h, when embryos are expected in the uterine horn (39), we observed that a third of the XER mice displayed embryos in the oviduct only, a third of the mice displayed embryos in both the oviduct and the uterus, and a third of the mice displayed embryos in the uterus alone (Supp. Fig. 6A). When we evaluate a day later at GD4 1200h, 4/7 XER mice displayed embryos in their uterus and the average number of embryos per mouse was lower than controls (Supp. Fig. 6B). At GD4 1200h, implantation sites were clearly observed in control mice using the blue dye reaction (40) (Fig. 3A). XER mice failed to display any implantation at all, and LER mice displayed faint implantation sites reminiscent of delayed implantation (41) (Fig 3A, B, C). When analyzed a day later for decidual sites at GD5 1200h, control and LER mice displayed visible deciduae whereas XER mice failed to show decidualization (Fig 3A, C). Since LERs exhibited a delayed implantation, we tested if the embryos displayed normal development at mid-gestation and if LER moms could deliver pups. LER moms had apparently normal embryos at GD13 1200h (Fig. 3B, D) and displayed normal number of pups at birth (Fig. 3E). Thus, despite loss of ESR1 in 90% of their glandular epithelium in the pre-implantation stage, LER mice can carry pregnancy to term.

**Figure 3.**
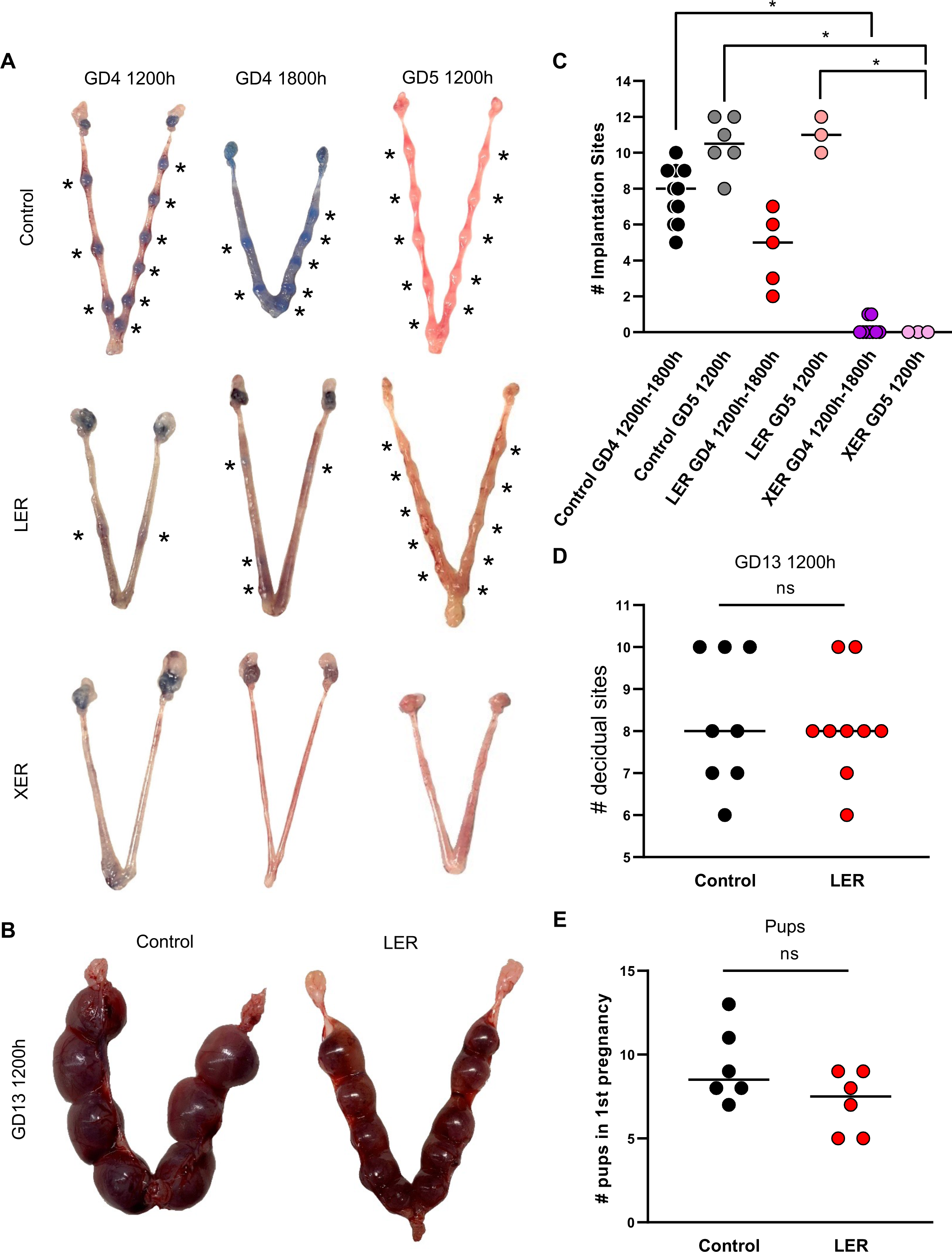
Implantation and pregnancy phenotypes of mice with epithelial ESR1 deletion. (A) Dissected uteri from control, LER, and XER mice at GD4 1200h, GD4 1800h, and GD5 1200h. Uteri at GD4 1200h and GD4 1800h were injected with blue dye prior to dissection for visualization of implantation sites. Asterisks indicate implantation/decidual sites. XER mice exhibit failed implantation while LER mice exhibit delayed implantation. (B) Dissected uteri from control and LER mice at GD13 1200h showing that LER mice display pregnancy progression comparable to controls. (C-D) Quantification of number of implantation sites at GD4 1200h-1800h and GD5 1200h in control, LER, and XER mice (C) and at GD13 1200h in control and LER mice (D). Each dot represents one uterus analyzed. Median values shown. Data analyzed using Mann-Whitney test. (E) Quantification of number of live pups born in first pregnancies of control and LER mice. Each dot represents one uterus analyzed. Median values shown. Data analyzed using Mann-Whitney test. (*) = *p<0.05*. (ns) = *p>0.05*.

### Mice with branchless glands fail to form an implantation chamber and display aberrant embryo-uterine axes alignment

Previous research from our lab has demonstrated that formation of an embryo implantation chamber is necessary for the alignment of the embryonic and the uterine axis, such that the inner cell mass of the blastocyst faces the mesometrial pole of the uterus in a V-shaped implantation chamber (42). This event coincides with a shift of PTGS2 expression from the luminal epithelium to the stroma underlying the implantation chamber (42). When we evaluate ESR1-deficient uteri at GD4 1200h, we observed that both XERs and LERs show a failure of: a) V-shaped implantation chamber formation, b) embryo-uterine axes alignment, and c) PTGS2 expression in the stroma under the embryo attachment site (Fig. 4A a-c”). Control embryos displayed an array of axis angle values with a median value of 24.30° (Fig. 4C), while embryos of LER and XER mice were misaligned with the ICM pointing in a perpendicular direction (median angle LER 87.70 ° and XER 88.6°) with respect to the M-AM axis (Fig. 4C). However, at GD4 1800h, LERs begin to display a small V-shaped chamber, improved alignment of the embryo-uterine axis (median angle 57.70 °), and PTGS2 expression in the stroma underlying the chamber (Fig. 4A d-e”, 4C). Further, by GD5 1200h, both control and LER uteri displayed comparable embryo development commensurate with epiblast stage (Fig. 4B). We failed to observe live embryos in XER mice at GD4 1800h (one misaligned embryo was observed in 1/10 uterine horns from five mice analyzed), suggesting that in the absence of branched glands and ESR1, attachment fails at GD4 1200h and embryos fail to survive beyond this stage.

**Figure 4.**
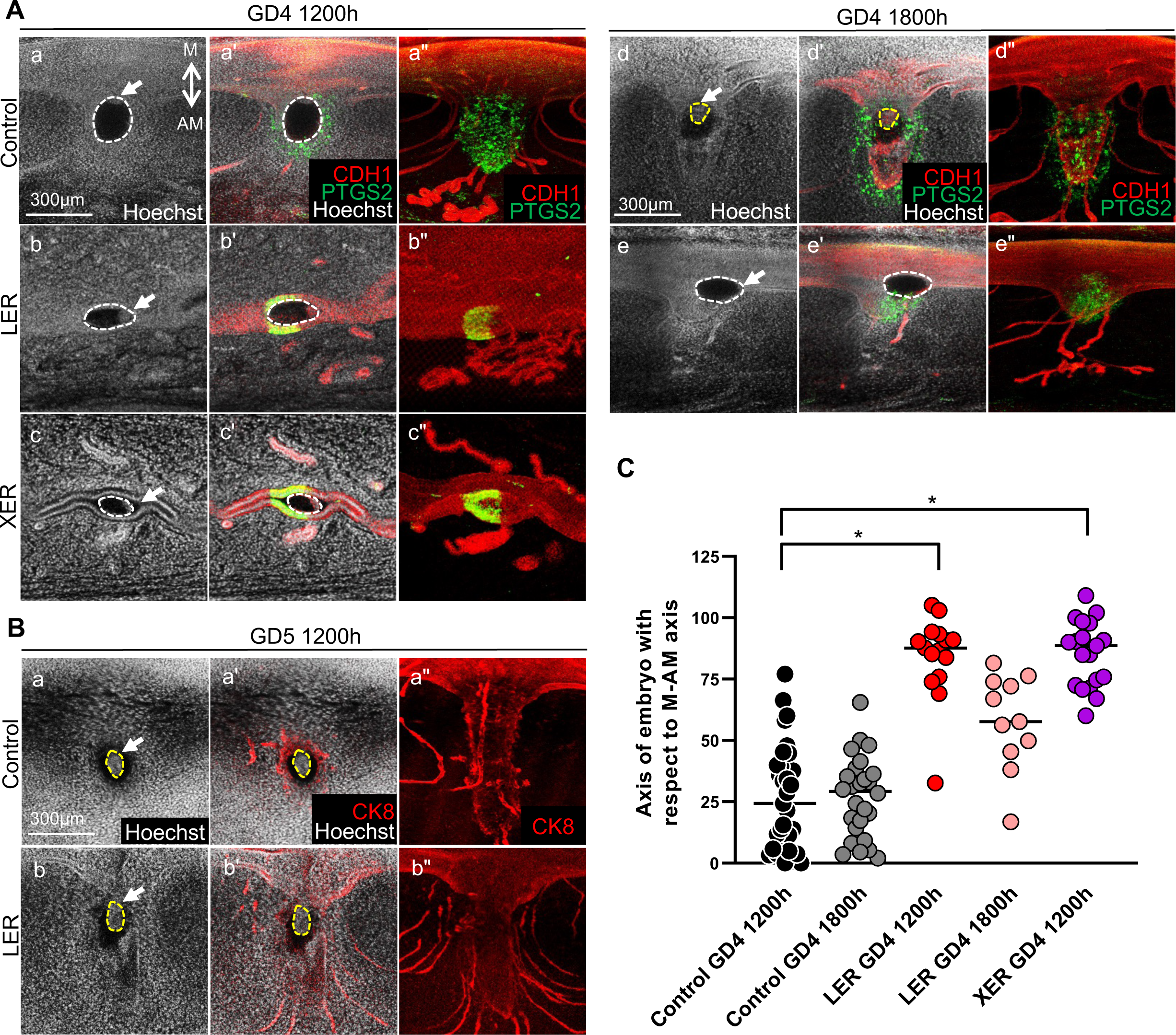
Embryo-uterine axes alignment phenotypes in mice with uterine epithelial ESR1 deletion. (A-B) Immunofluorescent images of embryos at implantation sites in control, LER, and XER mice at GD4 1200h, GD4 1800h, and GD5 1200h with staining for (A) Hoechst (nuclei, white), E-Cadherin (epithelium, red), and PTGS2 (green) or (B) Hoechst (nuclei, white) and Cytokeratin 8 (epithelium, red). Cross-sections were obtained from whole tissue confocal imaging. (") column images represent pseudo-3D cross-sections that are ∼150-300μm in thickness. All other images are 7μm in thickness. Mesometrial-antimesometrial (M-AM) axis is indicated (a), and white arrows point to the inner cell mass of embryos. White dotted lines represent the boundary of an embryo. Yellow dotted lines indicate the boundary of the epiblast. Scale bar, 300μm. (n=3) mice per genotype per stage. (C) Embryo axis angle measurements of control, LER, and XER mice at GD4 1200h and GD4 1800h. Angle measurements taken with respect to the M-AM axis. Each dot represents one embryo. Median values shown. Data analyzed using Kruskal-Wallis test with Dunn’s multiple comparisons. (*) = *p<0.05*.

### ESR1-deficient unbranched glands fail to express preimplantation *Lif*

Since ESR1-deficient mouse models have shown reduced LIF expression (11) we determined *Lif* mRNA expression using in situ hybridization. First, we observed that in control mice at GD0 1200h *Lif* is highly expressed in the luminal compartment but not in the glandular compartment (Supp Fig. 7). At pre-implantation stage GD3 1200h in controls, strong *Lif* expression was observed colocalizing with FOXA2 expressing uterine glands (Fig. 5A, Supp Fig. 7) and *Lif* expression was not observed in the luminal compartment. XER glands failed to express *Lif* while surprisingly, a small amount of *Lif* was observed in LER glands (Fig. 5A, B). Since we observed delayed implantation in LERs, we tested for *Lif* expression a few hours later at GD3 1800h. We discovered that while XER glands again failed to express *Lif*, LER glands showed an increase in *Lif* expression compared to GD3 1200h (Fig. 5A, B). We quantified total *Lif* volume normalized for gland volume using image segmentation protocols. Control glands express a uniformly high volume of *Lif* normalized for gland volume at GD3 1200h and GD3 1800h while LER and XER mice exhibited much lower volume of *Lif* at GD3 1200h. We observed a slight but significant increase in the amount of *Lif* volume in LER mice at GD3 1800h. Visually, we observed that although LER gland cells do not uniformly express *Lif,* a few gland cells express exaggerated amounts of *Lif* which is sufficient to support implantation.

**Figure 5.**
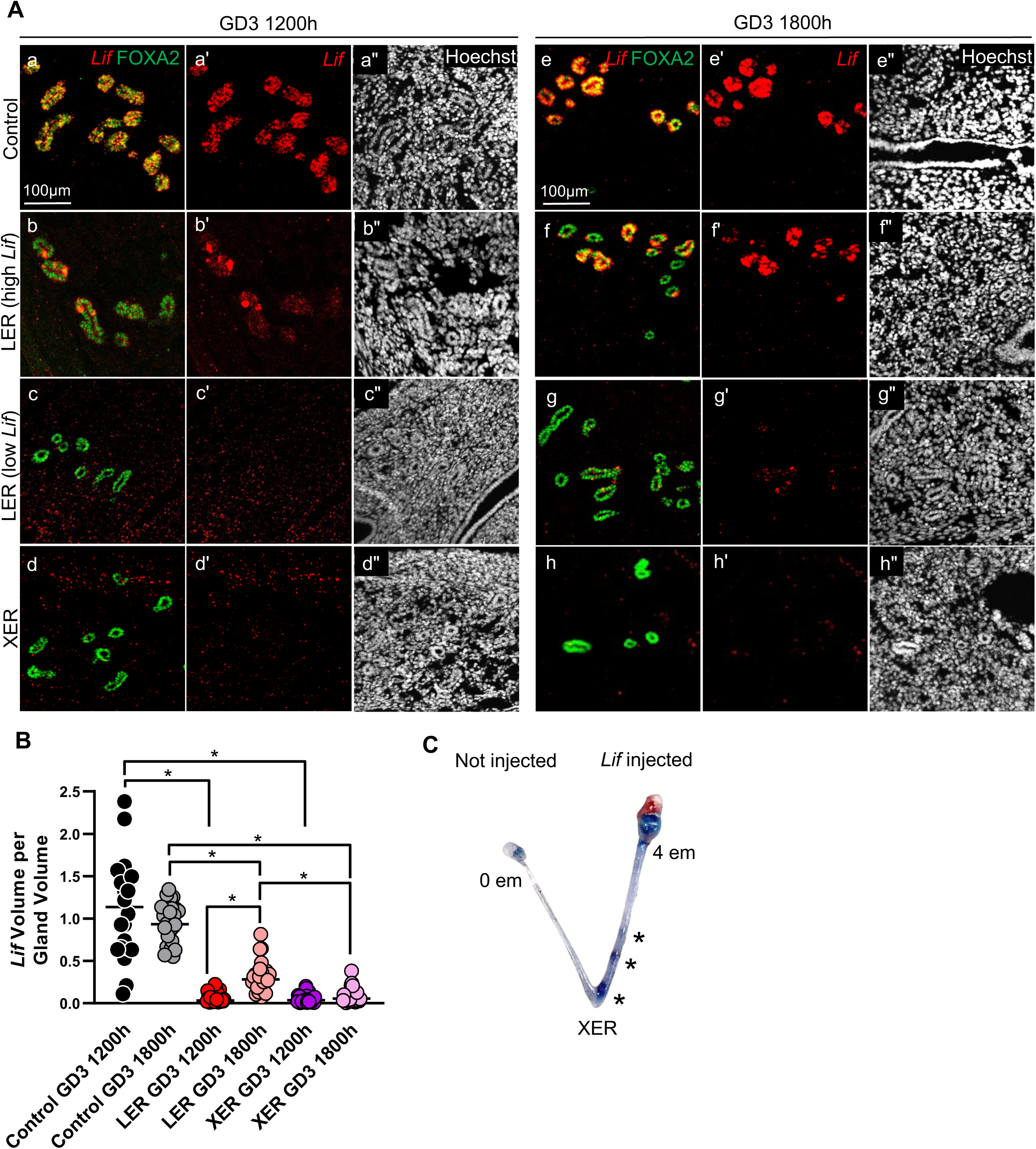
Epithelial ESR1 deletion mice display defects in Lif expression. (A) Uterine sections of control, LER, and XER mice at GD3 1200h and GD3 1800h with staining for Hoechst (nuclei, white), FOXA2 (gland marker, green), and *Lif* mRNA (red). *Lif* expression in LER glands is high in a subset of cells (b-b”, g-g”) and lower in others (c-c”, g-g”). Lif expression increases from GD3 1200h to GD3 1800h in LER glands while XER glands express undetectable levels of *Lif* at both time points. Scale bar, 100μm. (B) Quantitative analysis of the amount of *Lif* per gland volume in control, XER and LER mice at GD3 1200h and GD3 1800h. n=9 regions each from 3 mice per genotype per stage were analyzed. Data in (B) analyzed using Kruskal-Wallis test with Dunn’s multiple comparisons. (*) = *p<0.05*. (C) Uterine horn of XER mouse intraluminally injected with 1μg recombinant *Lif* into the right horn on GD3 1200h and dissected on GD4 1200h following blue dye injection. Asterisks indicate implantation rescue sites, and the total number of embryos found in each horn are shown.

### LIF supplementation partially rescues implantation in ESR-1 deficient uteri

Because we observed almost no *Lif* expression in XERs, we assessed whether recombinant LIF administration at GD3 could rescue implantation in XER mice at GD4 1200h. For XER uteri that did contain embryos, using both intraperitoneal and intraluminal routes of LIF administration (21, 22), we achieved 75% and 67% of implantation rescue, respectively (Fig. 5C, Supp. Fig. 8). Altogether, this demonstrates that the critical difference between XER mice with unbranched glands and LER mice with branched glands is the presence of a threshold amount of LIF in LER mice.

### Unbranched glands with E2-ESR1 signaling fail to express *Lif*

XER glands are unbranched but also lack ESR1 signaling. To separate the contributions of gland branching and ESR1 signaling for glandular *Lif* expression, we used pre-pubertal P21 mice that have unbranched glands and treated them with either E2 alone, E2+P4, or vehicle (sesame oil) to evaluate the effect of ovarian steroids on *Lif* expression. We observed that P21 glandular and luminal epithelium and the stroma express high levels of ESR1 in the presence of vehicle alone whereas ESR1 levels are reduced in the epithelium in the presence of two doses of E2 (Fig. 6A, a-a”, b-b”). On the other hand, ESR1 is highly expressed in glandular epithelium when P21 mice are treated with E2 and then P4 (Fig. 6A, c-c”). We also observed that P21 glands treated with vehicle or hormones express *Lif* in their luminal epithelium (Fig. 6 d-d”, e-e”, f-f”, 6C). Finally, although we observed varying levels of ESR1 expression in the glandular epithelium of mice treated with vehicle or hormones, E2 treatment failed to elicit *Lif* expression in glandular epithelium (Fig. 6A, B). Because it has been suggested that glandular *Lif* expression occurs under P4 dominant conditions (43) we also evaluated P21 mice treated with both E2 and P4 but again did not detect any *Lif* expression in the glands (Fig. 6A, f-f”). When we evaluated the 3D gland structure of the P21 uteri treated with different hormonal regimens (Fig. 6D), we observed that treatment with E2 alone or E2+P4, did not result in a change in gland length compared to vehicle treatment (Fig. 6E). Further, in all treatment groups glands were found to be primarily unbranched (Fig. 6F) and no glands with >3 branches were observed. Since ESR1 was expressed in the glandular compartment, the observed lack of *Lif* expression at P21 cannot be due to an absence of ESR1 signaling. Thus, branchless glands even in the presence of E2-ESR1 signaling fail to produce *Lif*.

**Figure 6.**
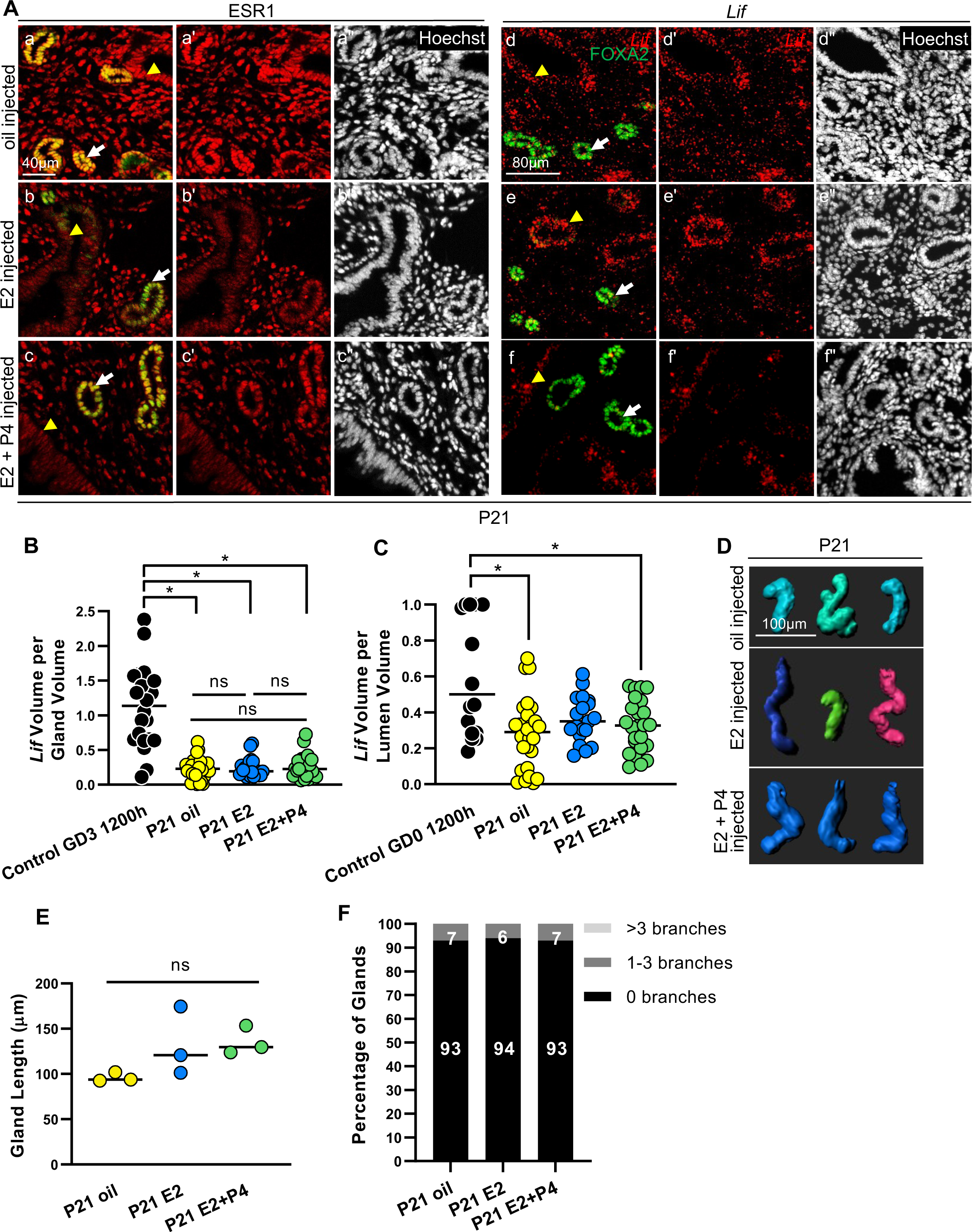
Pre-pubertal, unbranched glands fail to produce Lif even when ESR1 signaling is intact. (A) Uterine sections of P21 control mice injected with either oil, E2, or E2+P4 and stained for Hoechst (nuclei, white), FOXA2 (gland marker, green), ESR1 (a, b, c, red), and *Lif* mRNA (d, e, f, red). Scale bar, 40μm (a, b, c). Scale bar, 80μm (d, e, f). n=9 regions each from 3 mice per treatment. Quantitative analysis of (B) the amount of *Lif* normalized for gland volume and (C) the amount of *Lif* normalized for lumen volume. n=9 regions each from 3 mice per treatment. (B-C) Data analyzed using Mann-Whitney test. (*) = *p<0.05.* Yellow arrowheads point towards the luminal epithelial cells and white arrows points towards the glandular epithelial cells (D) Representative 3D reconstructions of P21 control uterine glands with various treatments. n=3 mice per treatment group. Scale bar, 100μm. Quantitative analyses of (E) average gland length measurements (each dot represents one mouse) and (F) percentage of glands with 0, 1-3, and >3 branches in P21 control mice with various treatments. Hormonal treatment does not affect gland length or gland branching. 85-450 glands analyzed per mouse. (E) Data analyzed using Mann-Whitney test. (*) = *p<0.05*. Two-proportion Z-test determined that the differences in percentage of gland branches between oil, E2, and E2+P4 treatment is not statistically significant.

## DISCUSSION

Uterine glands are exocrine glands with a distinct structure that play vital roles in implantation and pregnancy success. Exocrine gland structure is well known to contribute to gland function, however such structure-function relationship for uterine glands have not been described. In our study we discovered that developmental ESR1 signaling in different compartments of the uterus is necessary to build a branched gland. Further we determine that both branched structure of a uterine gland and ESR1 signaling are critical to preimplantation LIF production for gland function in implantation.

### Secretory epithelial domains in branched uterine glands

During development, mouse uterine glands bud off the uterine lumen at postnatal day (PD) 4. They elongate into the stroma, going through changes in morphology including tear drop, elongated, and sinuous stages until some begin branching at P21 (3). In the adult mouse, uterine glands are prominently branched (36). Human uterine glands in the functionalis and the basalis layer have also been characterized for 3D structure (45). Glands in the functionalis layer are coiled and show some branching while glands in the basalis layer show more prominent branching compared to the functionalis glands. Exocrine glands such as uterine glands are generally classified as tubular or branched. Tubular exocrine glands do not have designated secretory cells, although tubular sweat glands have a coiled portion that holds secretions (46). In contrast to tubular glands, branched exocrine glands have acinar end-pieces that are secretory in nature (47). The presence of functional sub-structures in uterine glands (end-pieces, ducts, coils) has not been established. It is possible that the individual uterine gland has a ductal portion and a secretory end-piece (the branched end), however it is also possible that individual uterine glands are end-pieces, and the uterine lumen accumulates the secretions as ducts do in other branched exocrine glands. Our study establishes that in mice, unbranched uterine glands (XERs and P21) are unable to produce *Lif.* This supports the idea that branched portions of uterine glands may carry the secretory function. Whether this is true for human uterine glands will be an avenue for future investigatioin.

### ESR1 signaling and gland structure in mammary and uterine glands

E2 signaling plays a key role in establishing the branching pattern of the exocrine mammary glands that are essential for lactation (29, 30). A whole body deletion of *Esr1* (*Esr1KO*) leads to mammary glands with minimal branching (28, 29), while an epithelium-specific deletion of *Esr1* displays a loss of duct elongation and side branching (30). Thus, gland proliferation and branching are ESR1 signaling-dependent in the mammary gland. In mice with continuous estrous, the uterine glands display a cribiform structure (48), suggesting that aberrant ESR1 signaling must impact uterine gland shape. In the uterine glands we determined that neonatal deletion of ESR1 in the epithelial and stromal compartments resulted in gland buds while deletion only in the epithelial compartment resulted in glands that elongate but fail to branch. Together this suggests that stromal ESR1 signaling regulates gland bud outgrowth and elongation in a non-cell-autonomous manner while epithelial ESR1 signaling regulates gland branching during the formative phases of the uterine glands. The gland bud phenotype is remarkably similar to gland structure observed in uteri with neonatal deletion of *Foxa2 (*Fig. 4F in reference (49)). Depletion of ESR1 in the adult epithelium does not cause any quantifiable structural gland phenotypes, suggesting that epithelial ESR1 signaling during puberty is required for gland branching. Once glands are branched, reduced E2-ESR1 signaling did not affect gland structure during estrous cycles. This is in contrast with loss of FOXA2 in the adult epithelium, which results in loss of *Lif* and the gland structure appears abnormal (Fig 4F in reference (49)). We propose that ESR1 signaling and FOXA2 may work together for gland bud elongation and for *Lif* production, however they may be redundant to maintain branched glands post-puberty.

### Utility of Pax2 Cre for studying uterine epithelium

*Wnt7a^Cre^ ESR1^flox/flox^*mice do not support embryo entry into the uterus as the embryos die in the oviduct precluding analysis of embryo implantation studies. Using the *Pax2^Cre^* line we were able to bypass the oviductal phenotype and embryos were observed in the uterus during the pre-implantation and implantation time points. This may be because we have incomplete and patchy expression of the *Pax2^Cre^* lineage in the oviductal epithelium. Recently, Hancock et. al (50) determined that *Wnt7a^Cre^ ESR1^flox/flox^* mice also display ectopic expression of gland marker FOXA2 in the uterine luminal epithelium at GD3 1200h. In *Wnt7a^Cre^* mouse, *Wnt7a* coding exons are replaced by *Cre* making this model a heterozygous for *Wnt7a* gene (10). It is unknown if the ectopic expression of FOXA2 in the *Wnt7a^Cre^ ESR1^flox/flox^* lumen is due to depletion of ESR1 alone or due to the additional loss of one copy of *Wnt7a*. WNT7A has key functions in uterine epithelium and gland specification (20, 51) and could interfere with the ESR1 mutant analysis. We did not observe ectopic expression of FOXA2 in our *Pax2^Cre^ ESR1^flox/flox^* uterine lumen and we observed severe gland branching phenotypes in both models where ESR1 signaling is disrupted in the embryonic epithelium. This supports the idea that alternate Cre lines such as *Pax2^Cre^*may be powerful tools to study the impact of loss of gene function in the embryonic uterine epithelium where *Wnt7a^Cre^* cannot be used.

### E2-ESR1 signaling and LIF production

LIF is an essential gland secretion that is absolutely necessary for implantation. *Lif* knockout mice and other genetic models that fail to produce LIF exhibit embryo implantation failure. ESR1 signaling is predicted to be upstream of *Lif* expression because ESR1 binding sites are observed in the coding region and in the 3’UTR of the *Lif* gene (52). In uterine biology literature, often a bolus of E2 injection results in increased levels of *Lif* mRNA which is usually observed using whole tissue analysis such as qPCR. Since E2-ESR1 signaling can upregulate *Lif* in both luminal and glandular epithelium but only glandular LIF promotes embryo implantation, it is necessary that these studies be interpreted carefully and the location of *Lif* upregulation be determined when employing E2 treatment. We observed that ovulatory E2 only induces high *Lif* expression in the luminal epithelium, while in P4-dominant pre-implantation stages, *Lif* is confined to the glandular epithelium. It has been shown in various mammalian species such as mice (43), hamsters (53), marmosets (54), and rabbits (55) that E2 alone is unable to induce *Lif* expression in the glands and other factors may be necessary for LIF production. Our work suggests that in addition to E2-ESR1 signaling, branched structure of the uterine gland is necessary for *Lif* synthesis. *Lif* mRNA was negligible when uterine glands lacked ESR1 signaling in addition to a branched structure or in the case of prepubertal glands where although glands had E2-ESR1 signaling, glands were unbranched. Our quantitative data from both mouse models with epithelial deletion of ESR1 suggests that glands with greater than three branches are key for producing threshold levels of glandular LIF to support implantation resulting in pregnancy success. We also observed that uteri with normally branched glands but severely reduced ESR1 signaling were still able to produce threshold levels of LIF to support embryo implantation and pregnancy success. This indicates that gland functional ability (LIF secretion and implantation) is dependent on its structure, particularly whether a gland is branched or unbranched.

### Uterine gland structure – function relationships

Loss of ESR1 signaling in the mammary glands results in poorly developed branched structure and compromises the lactation function of the glands. We also show that unbranched glands compromise *Lif* secretion, resulting in implantation failure, and this phenotype is partially rescued when pregnancy is supplemented with LIF. It is critical to note that supplemental LIF can rescue both implantation and pregnancy, at least partially, in genetic mutants where glands are branched but simply fail to produce LIF (adult epithelial ESR1 deletion, this study, and adult epithelial FOXA2 deletion (22)). However, if a genetic mutant displays failure for glands to branch and fails to produce *Lif*, then implantation is rescued (embryonic ESR1 deletion, this study and neonatal FOXA2 deletion (22, 49, 56)), but pregnancy fails to continue post-implantation (22). These data support essential functions for branched uterine glands in the post-implantation phase.

Intriguingly, mice with neonatal or adult epithelial deletion of FOXA2 displayed implantation failure, but the embryos apparently entered a diapause state. Embryos in our embryonic epithelial deletion of ESR1 did not survive beyond implantation stages. This could be due to differences in the progesterone signaling in these mutant mice and will be a subject of future studies. However, even with embryos that entered diapause in FOXA2 deletion mutants, viability of embryos that failed to implant was higher if glands were branched compared to bud staged glands (49), supporting multiple roles for branched uterine glands beyond LIF secretion and embryo implantation.

From our research into the structure-function relationship of uterine glands in the context of pregnancy outcome, thus far we have gathered new insights regarding how uterine estrogen signaling affects gland development and gland shape, and how gland structure contributes to the induction of *Lif* and events surrounding embryo implantation. To expand the knowledge of how branched and unbranched glands differ in functional capability, further research is needed to determine the secretory factors a branched gland makes and if there are specific secretory cell types in uterine glands similar to mammary and salivary glands (4). Additionally, it will be useful to assess the cellular differentiation events that define how glands bud off the main luminal epithelial tube and the steroid hormone dependent and independent signaling mechanisms that guide gland bud elongation and branching. Further, post-pubertal gland development especially in the context of regeneration in the post menses and the post-partum uterus remain understudied. More research is therefore needed to characterize how cyclicity and pregnancy remodel uterine glands and whether and how new gland buds may arise in adulthood. Gland structure-function dynamics in humans are challenging since it will be necessary to assess both the basalis and the functionalis layers of endometrial glands.

Therefore, the mouse can provide novel insights into gland structure-function relationships relevant to early pregnancy in mammals.

## AUTHOR CONTRIBUTIONS

K.G, S.F. and R.A. designed the experiments; K.G, S.F., M.S., J.L., A.B., and J.H. performed experiments; K.G, S.F., M.S., J.L., and R.A. analyzed the data. K.G., S.F., X.Y., and R.A. interpreted the results; K.G., S.F., and R.A. wrote the manuscript.

## ACKNOWLEDGEMENTS

We acknowledge Sameed Khan for contributing to image analysis discussions and Frank Lawrence at the Center for Statistical Training and Consulting at Michigan State University for help with statistical analysis. We are also grateful to Drs. Asgerally Fazleabas, Nataki Douglas and Gregory Burns for critical analysis and research discussions.

## DATA AVAILABILITY STATEMENT

Data generated in the manuscript is available upon reasonable request to the corresponding author.

---

**Supp. Figure 1.**
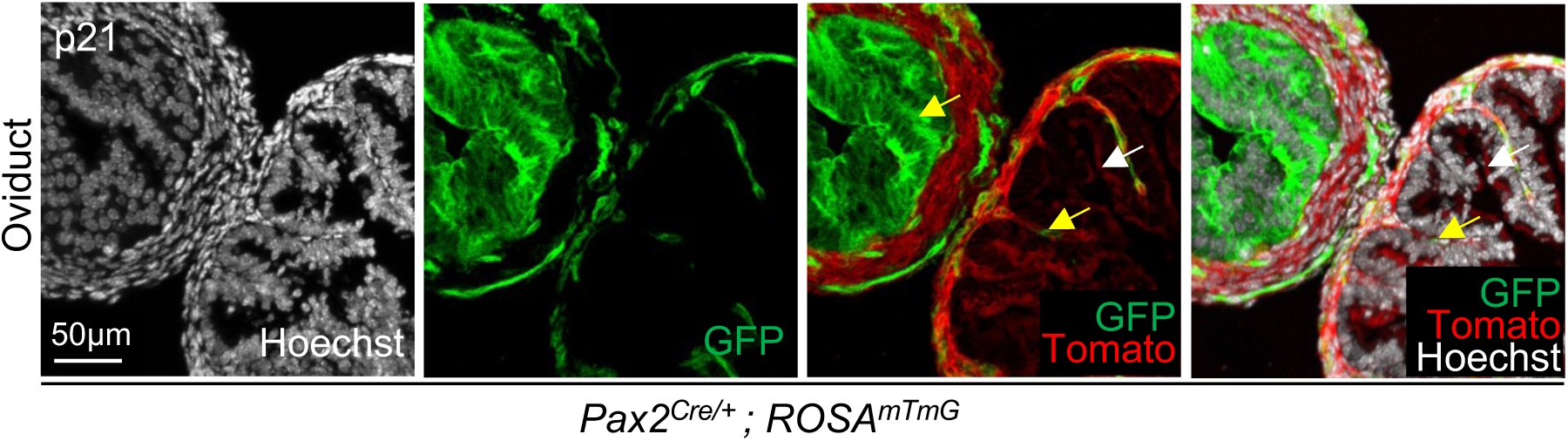
Pax2 lineage in oviductal epithelium. (A) *Pax2^Cre/+^* ; *ROSA^mT/mG^* mouse oviduct with Hoechst (nuclei, white), GFP (green), and Tomato (red) demonstrating patchy Pax2 lineage GFP reporter expression in the oviductal epithelium. Scale bar, 50μm. Yellow arrows indicate GFP lineage expression in oviductal epithelium. White arrows indicate Tomato expression and absence of GFP expression in oviductal epithelium.

**Supp. Figure 2.**
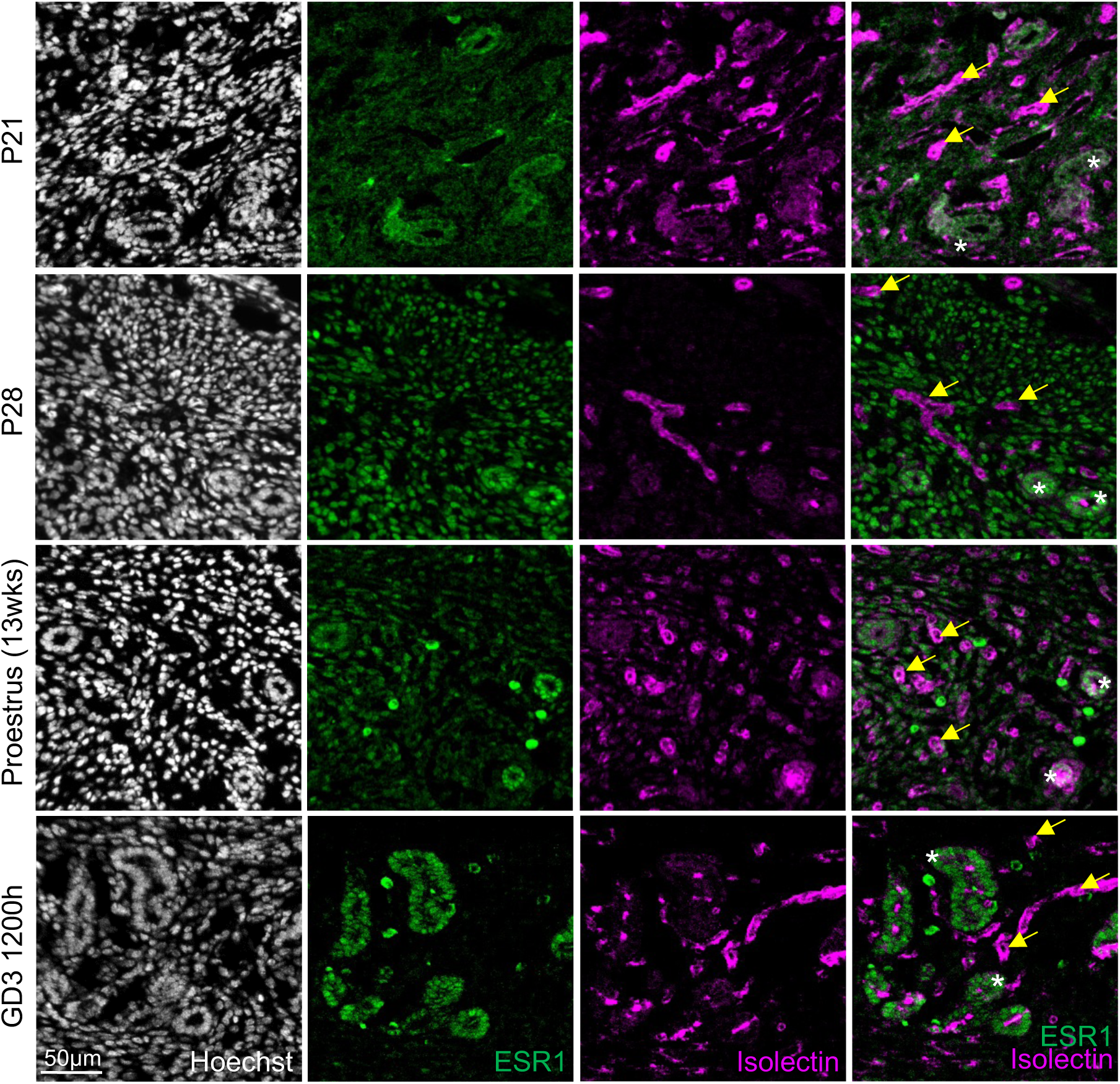
ESR1 is not expressed in uterine vasculature. Uterine sections from control mice stained with Hoechst (nuclei, white), ESR1 (green), and Isolectin (endothelial marker, magenta). Scale bar, 50μm. P21, n=2 mice; P28, Proestrus and GD3 1200h n=3 mice per age/stage. Yellow arrows indicate regions of vasculature and white asterisks indicate cells expressing ESR1. No overlap is observed between the two.

**Supp. Figure 3.**
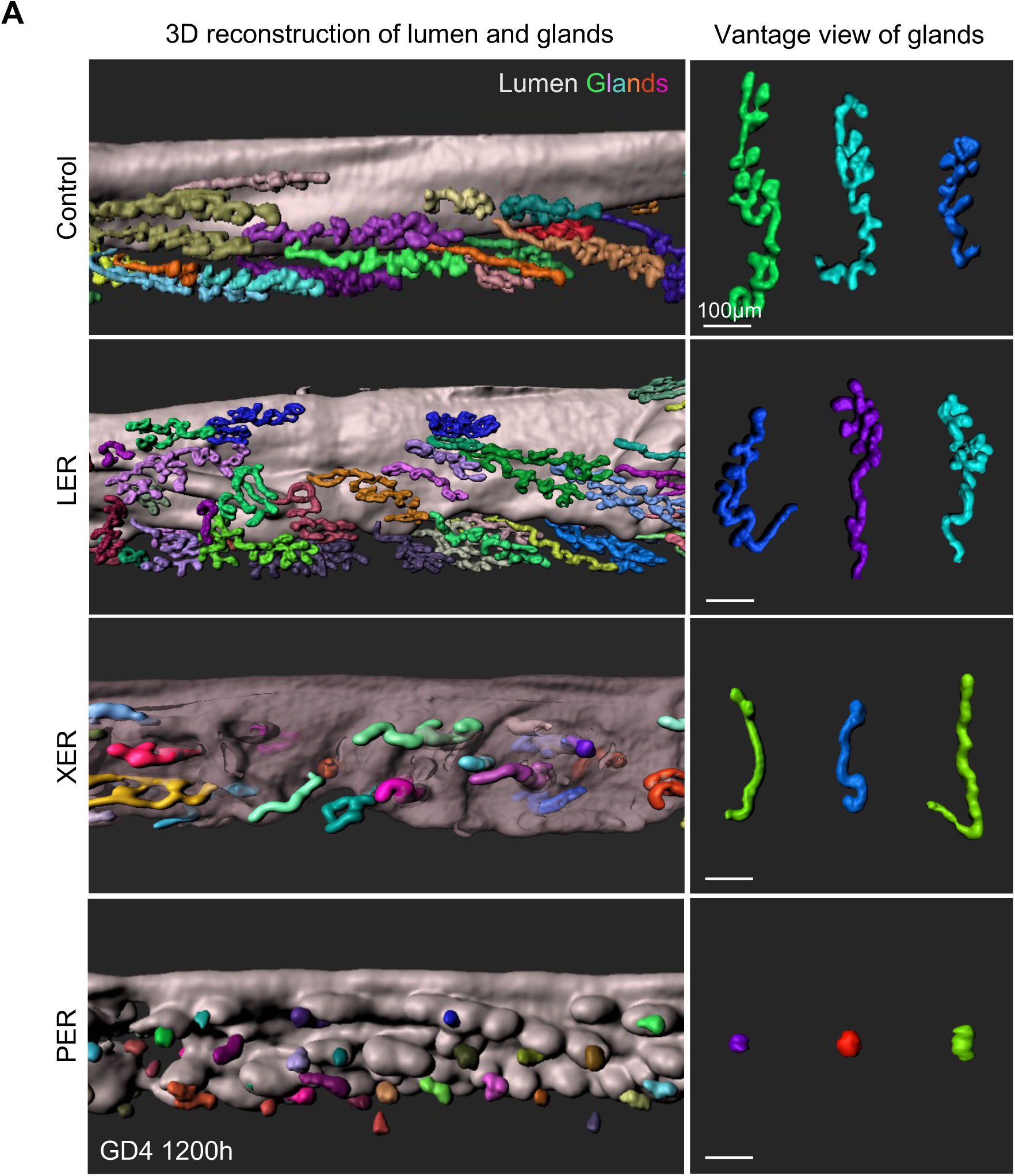
3D reconstructions of uterine lumen and glands. (A) Whole uterine tissue was stained with CDH1 (epithelial marker) and FOXA2 (gland marker). Following confocal imaging at 10x magnification, 3D surfaces (left) of lumen and glands were made for visualization of structures. Vantage view (right) of glands allows for comparison of individual glands. Scale bar, 100μm.

**Supp. Figure 4.**
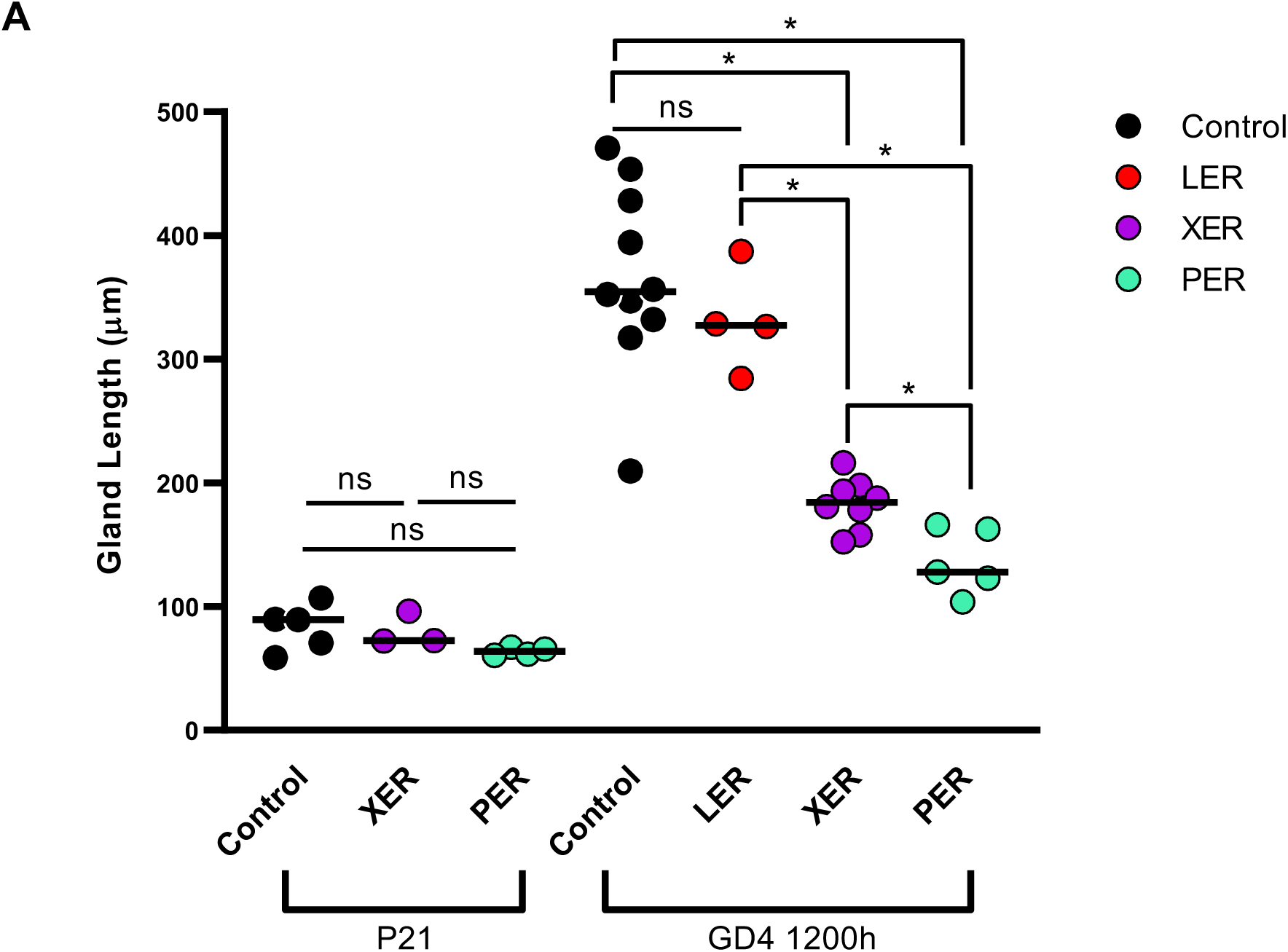
Neonatal and embryonic deletion results in reduced gland length during implantation. (A) Quantitative analysis of average gland length measurements at P21 and GD4 1200h. (n=3) mice/genotype/stage. 75-400 glands analyzed per mouse. Dot represents average gland length measurement from a single mouse. Data analyzed using Mann-Whitney test (*) = *p*<0.05. (ns) = *p*>0.05.

**Supp. Figure 5.**
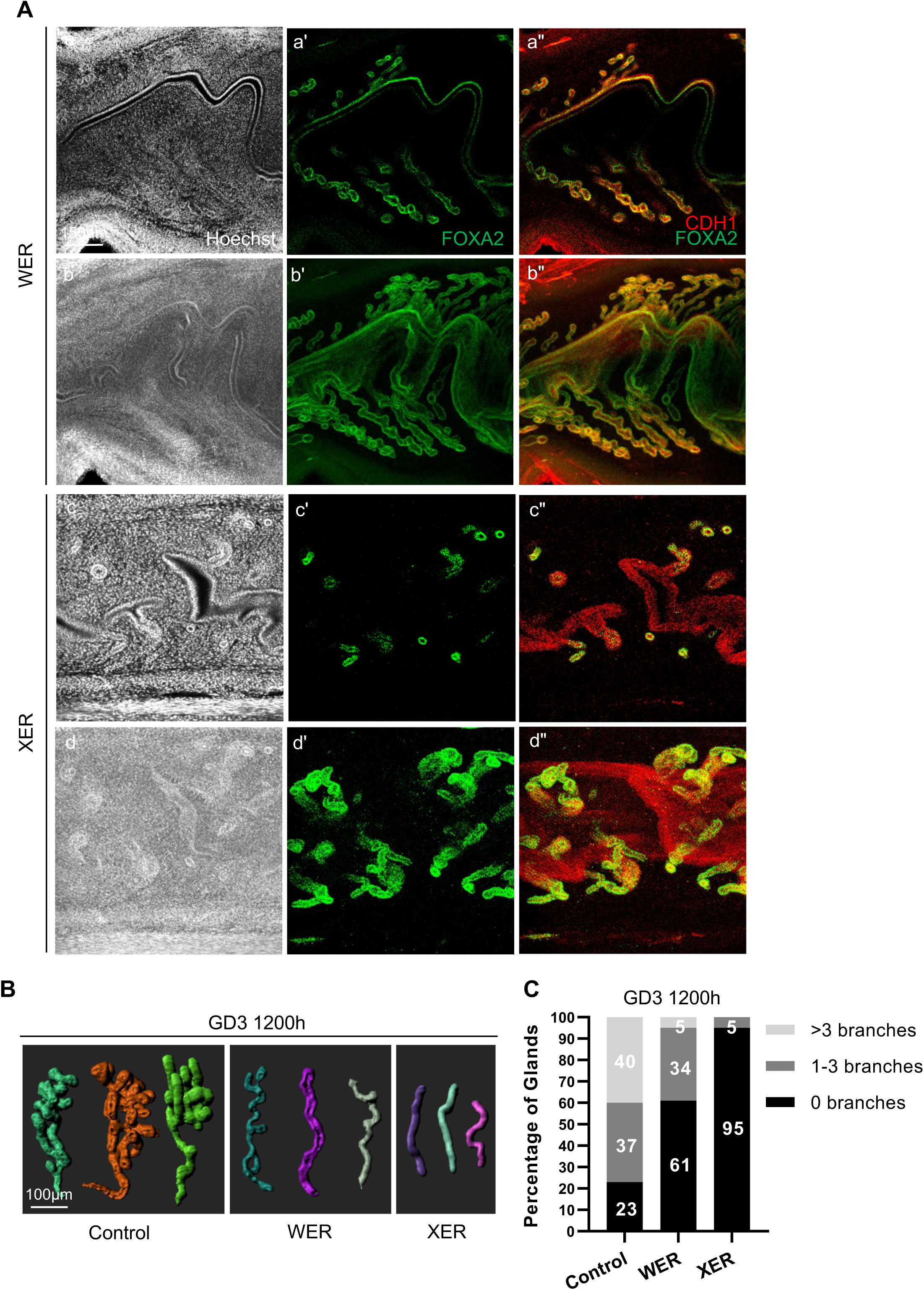
Gland branching is disrupted when two different Cre lines are used for embryonic epithelial Esr1 deletion. (A) Immunofluorescent images of (a-b) WER and (c-d) XER GD3 1200h uterine lumen and glands stained with Hoechst (nuclei, white), FOXA2 (gland marker, green), and CDH1 (epithelium, red). (a, a’, a”, c, c’, c”) Indicates 7μm slice view. (b, b’, b”, d, d’, d”) Indicates pseudo-3D view (150μm slice). Scale bar, 150μm. (B) Representative 3D reconstructions of control, WER, and XER glands previously stained with gland marker FOXA2 and imaged using a confocal microscope. Scale bar, 100μm. (n=3) mice. (C) Quantitative analysis of percentage of glands with 0, 1-3, and >3 branches in control, WER, and XER mice at GD3 1200h. Controls (n=3 mice, 6 uterine horns); WERs and XERs (n=2 mice, 4 uterine horns). 100-375 glands analyzed per mouse. Two-proportion Z-test determined that the differences in percentage of gland branches between control and WER glands, control and XER glands and WER and XER glands is statistically significant.

**Supp. Figure 6.**
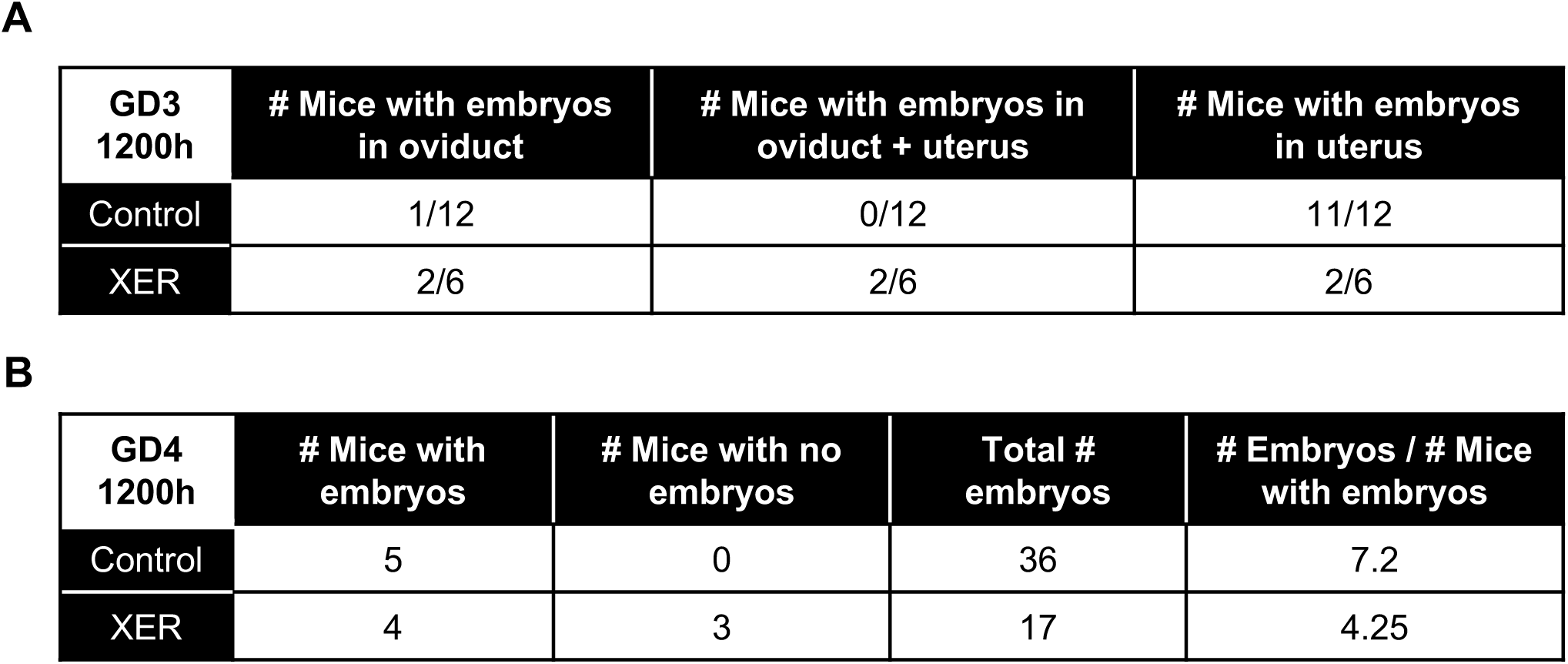
*Pax2^Cre^* deletion of ESR1 partially affects embryo entry into the uterus. (A) Table indicating location of embryos in control and XER mice at GD3 1200h when embryos should be in the uterus. (B) Table indicating number of control and XER mice with and without embryos at GD4 1200h along with the total number of embryos observed and the average number of embryos with respect to mice where embryos are present in the uterus

**Supp. Figure 7.**
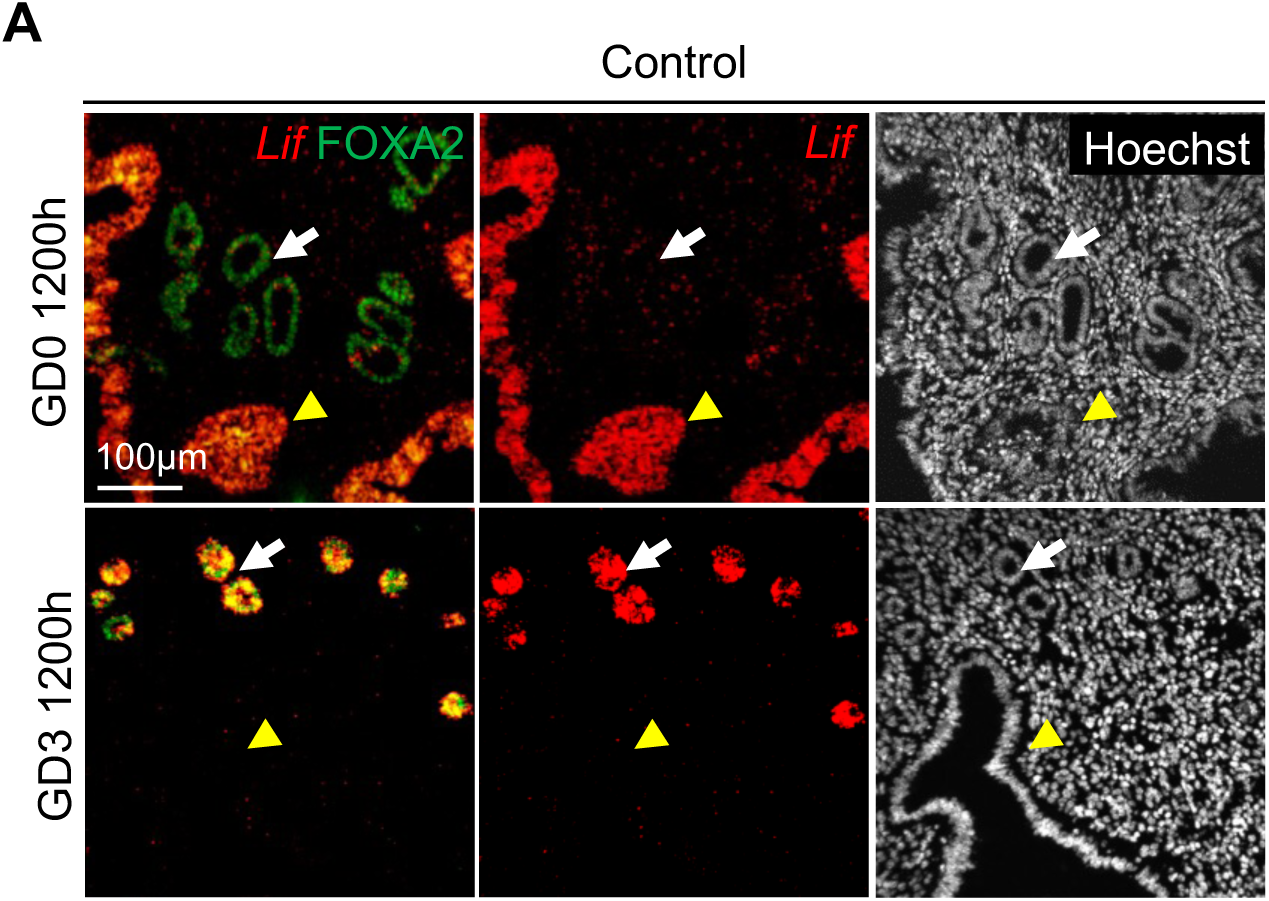
Differential Lif expression in luminal and glandular epithelium in early pregnancy. (A) Uterine section of control mouse at GD0 1200h and GD3 1200h with staining for Hoechst (nuclei, white), FOXA2 (gland marker, green), and *Lif* mRNA (red). *Lif* expression is evident in only the luminal epithelium at GD0 1200h and only the glandular epithelium at GD3 1200h. White arrows indicate glandular epithelium and yellow arrowheads indicate luminal epithelium. Scale bar, 100μm. (n=3) mice per stage.

**Supp. Figure 8.**
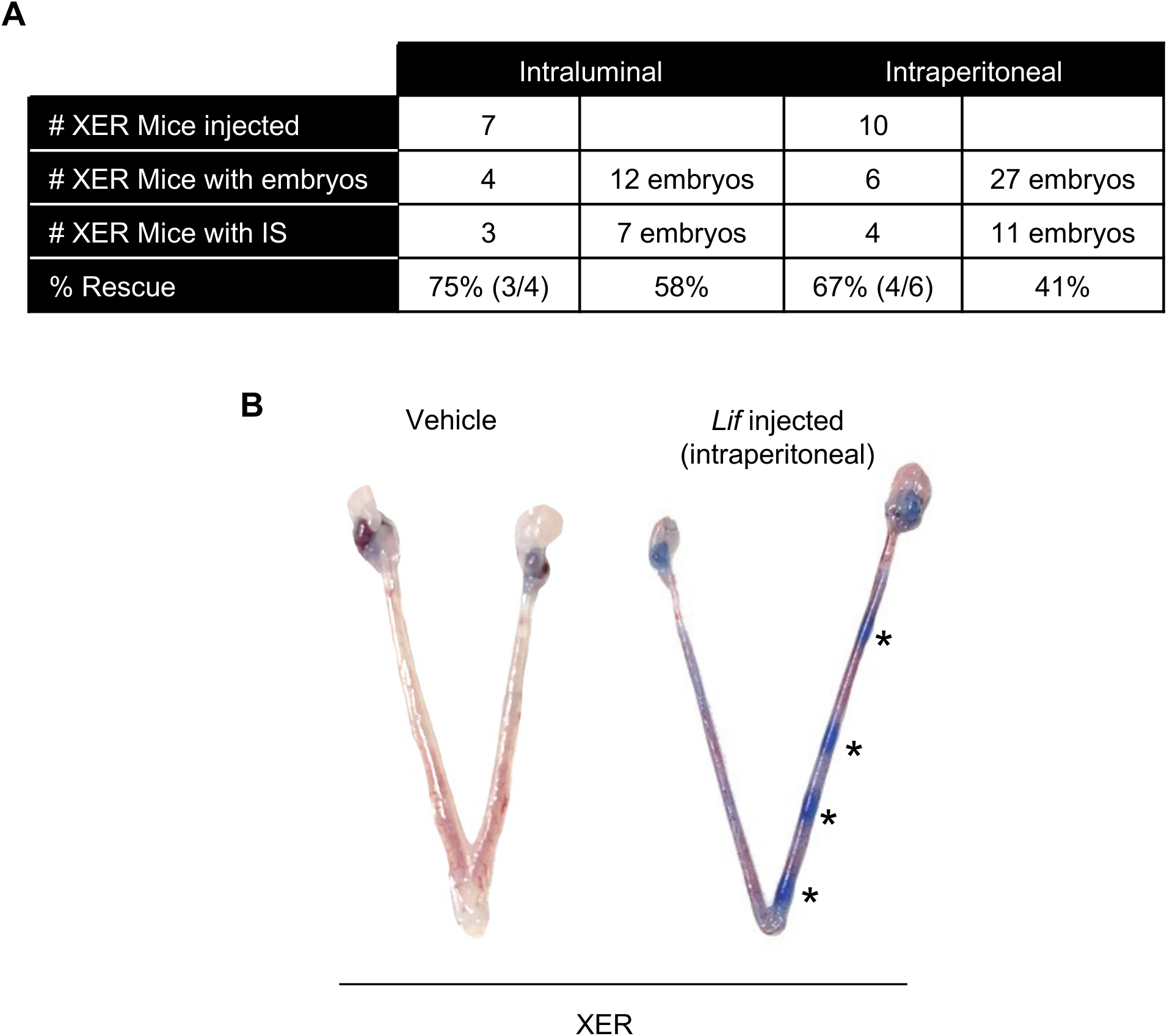
Implantation rescue in ESR-1 depleted mice with supplemental Lif. (A) Table indicates the total number of XER mice injected either intraluminally (1μg) or intraperitoneally (10μg) with *Lif*, the number mice where embryos were observed in the uterus, the number of mice with implantation sites, and the percent rescue of mice with implantation sites compared to mice with embryos. The total number of embryos observed in the uterus and the total number of implanted embryos along with the corresponding percent rescue is also indicated. (B) Dissected GD4 1200h uteri from XER mice injected intraperitoneally with either 10μg *Lif* or vehicle (PBS). Asterisks indicate blue dye sites.

## References

1. Kelleher AM, DeMayo FJ, and Spencer TE. Uterine Glands: Developmental Biology and Functional Roles in Pregnancy. Endocrine reviews. 2019;40(5):1424–45.

2. Gray CA, Bartol FF, Tarleton BJ, Wiley AA, Johnson GA, Bazer FW, et al. Developmental Biology of Uterine Glands1. Biology of reproduction. 2001;65(5):1311–23.

3. Vue Z, Gonzalez G, Stewart CA, Mehra S, and Behringer RR. Volumetric imaging of the developing prepubertal mouse uterine epithelium using light sheet microscopy. Molecular reproduction and development. 2018;85(5):397–405.

4. Khan S, Fitch S, Knox S, and Arora R. Exocrine gland structure-function relationships. *Development (Cambridge*, England). 2022;149(1).

5. Stewart CL, Kaspar P, Brunet LJ, Bhatt H, Gadi I, Köntgen F, et al. Blastocyst implantation depends on maternal expression of leukaemia inhibitory factor. Nature. 1992;359(6390):76-9.

6. Chen JR, Cheng JG, Shatzer T, Sewell L, Hernandez L, and Stewart CL. Leukemia inhibitory factor can substitute for nidatory estrogen and is essential to inducing a receptive uterus for implantation but is not essential for subsequent embryogenesis. Endocrinology. 2000;141(12):4365–72.

7. Cui J, Shen Y, and Li R. Estrogen synthesis and signaling pathways during aging: from periphery to brain. Trends in molecular medicine. 2013;19(3):197–209.

8. Curtis Hewitt S, Couse JF, and Korach KS. Estrogen receptor transcription and transactivation: Estrogen receptor knockout mice: what their phenotypes reveal about mechanisms of estrogen action. Breast cancer research : BCR. 2000;2(5):345–52.

9. Dupont S, Krust A, Gansmuller A, Dierich A, Chambon P, and Mark M. Effect of single and compound knockouts of estrogen receptors alpha (ERalpha) and beta (ERbeta) on mouse reproductive phenotypes. *Development (Cambridge*, England). 2000;127(19):4277–91.

10. Winuthayanon W, Hewitt SC, Orvis GD, Behringer RR, and Korach KS. Uterine epithelial estrogen receptor α is dispensable for proliferation but essential for complete biological and biochemical responses. Proceedings of the National Academy of Sciences of the United States of America. 2010;107(45):19272–7.

11. Pawar S, Laws MJ, Bagchi IC, and Bagchi MK. Uterine Epithelial Estrogen Receptor-α Controls Decidualization via a Paracrine Mechanism. *Molecular endocrinology (Baltimore*, Md). 2015;29(9):1362–74.

12. Winuthayanon W, Bernhardt ML, Padilla-Banks E, Myers PH, Edin ML, Lih FB, et al. Oviductal estrogen receptor α signaling prevents protease-mediated embryo death. eLife. 2015;4:e10453.

13. Tarleton BJ, Braden TD, Wiley AA, and Bartol FF. Estrogen-induced disruption of neonatal porcine uterine development alters adult uterine function. Biology of reproduction. 2003;68(4):1387–93.

14. Tarleton BJ, Wiley AA, and Bartol FF. Endometrial development and adenogenesis in the neonatal pig: effects of estradiol valerate and the antiestrogen ICI 182,780. Biology of reproduction. 1999;61(1):253–63.

15. Carpenter KD, Gray CA, Bryan TM, Welsh TH, Jr., and Spencer TE. Estrogen and antiestrogen effects on neonatal ovine uterine development. Biology of reproduction. 2003;69(2):708–17.

16. Ogasawara Y, Okamoto S, Kitamura Y, and Matsumoto K. Proliferative pattern of uterine cells from birth to adulthood in intact, neonatally castrated, and/or adrenalectomized mice, assayed by incorporation of [125I]iododeoxyuridine. Endocrinology. 1983;113(2):582–7.

17. Bigsby RM, and Cunha GR. Effects of progestins and glucocorticoids on deoxyribonucleic acid synthesis in the uterus of the neonatal mouse. Endocrinology. 1985;117(6):2520–6.

18. Jefferson WN, Padilla-Banks E, Suen AA, Royer LJ, Zeldin SM, Arora R, et al. Uterine Patterning, Endometrial Gland Development, and Implantation Failure in Mice Exposed Neonatally to Genistein. Environmental health perspectives. 2020;128(3):37001.

19. Filant J, and Spencer TE. Endometrial glands are essential for blastocyst implantation and decidualization in the mouse uterus. Biology of reproduction. 2013;88(4):93.

20. Dunlap KA, Filant J, Hayashi K, Rucker EB, 3rd, Song G, Deng JM, et al. Postnatal deletion of Wnt7a inhibits uterine gland morphogenesis and compromises adult fertility in mice. Biology of reproduction. 2011;85(2):386–96.

21. Jeong JW, Kwak I, Lee KY, Kim TH, Large MJ, Stewart CL, et al. Foxa2 is essential for mouse endometrial gland development and fertility. Biology of reproduction. 2010;83(3):396–403.

22. Kelleher AM, Peng W, Pru JK, Pru CA, DeMayo FJ, and Spencer TE. Forkhead box a2 (FOXA2) is essential for uterine function and fertility. Proceedings of the National Academy of Sciences of the United States of America. 2017;114(6):E1018–e26.

23. Filant J, Zhou H, and Spencer TE. Progesterone inhibits uterine gland development in the neonatal mouse uterus. Biology of reproduction. 2012;86(5):146, 1-9.

24. Rosario GX, and Stewart CL. The Multifaceted Actions of Leukaemia Inhibitory Factor in Mediating Uterine Receptivity and Embryo Implantation. American journal of reproductive immunology (New York, NY : 1989). 2016;75(3):246–55.

25. Shen MM, and Leder P. Leukemia inhibitory factor is expressed by the preimplantation uterus and selectively blocks primitive ectoderm formation in vitro. Proceedings of the National Academy of Sciences of the United States of America. 1992;89(17):8240–4.

26. Song H, Lim H, Das SK, Paria BC, and Dey SK. Dysregulation of EGF family of growth factors and COX-2 in the uterus during the preattachment and attachment reactions of the blastocyst with the luminal epithelium correlates with implantation failure in LIF-deficient mice. *Molecular endocrinology (Baltimore*, Md). 2000;14(8):1147–61.

27. Curtis Hewitt S, Goulding EH, Eddy EM, and Korach KS. Studies using the estrogen receptor alpha knockout uterus demonstrate that implantation but not decidualization-associated signaling is estrogen dependent. Biology of reproduction. 2002;67(4):1268–77.

28. Bocchinfuso WP, Lindzey JK, Hewitt SC, Clark JA, Myers PH, Cooper R, et al. Induction of mammary gland development in estrogen receptor-alpha knockout mice. Endocrinology. 2000;141(8):2982–94.

29. Sternlicht MD, Kouros-Mehr H, Lu P, and Werb Z. Hormonal and local control of mammary branching morphogenesis. Differentiation; research in biological diversity. 2006;74(7):365–81.

30. Feng Y, Manka D, Wagner KU, and Khan SA. Estrogen receptor-alpha expression in the mammary epithelium is required for ductal and alveolar morphogenesis in mice. Proceedings of the National Academy of Sciences of the United States of America. 2007;104(37):14718–23.

31. Gieske MC, Kim HJ, Legan SJ, Koo Y, Krust A, Chambon P, et al. Pituitary gonadotroph estrogen receptor-alpha is necessary for fertility in females. Endocrinology. 2008;149(1):20–7.

32. Soyal SM, Mukherjee A, Lee KY, Li J, Li H, DeMayo FJ, et al. Cre-mediated recombination in cell lineages that express the progesterone receptor. Genesis (New York, NY : 2000). 2005;41(2):58–66.

33. Ohyama T, and Groves AK. Generation of Pax2-Cre mice by modification of a Pax2 bacterial artificial chromosome. Genesis (New York, NY : 2000). 2004;38(4):195–9.

34. Daikoku T, Ogawa Y, Terakawa J, Ogawa A, DeFalco T, and Dey SK. Lactoferrin-iCre: a new mouse line to study uterine epithelial gene function. Endocrinology. 2014;155(7):2718–24.

35. Hewitt SC, Kissling GE, Fieselman KE, Jayes FL, Gerrish KE, and Korach KS. Biological and biochemical consequences of global deletion of exon 3 from the ER alpha gene. FASEB journal : official publication of the Federation of American Societies for Experimental Biology. 2010;24(12):4660–7.

36. Arora R, Fries A, Oelerich K, Marchuk K, Sabeur K, Giudice LC, et al. Insights from imaging the implanting embryo and the uterine environment in three dimensions. *Development (Cambridge*, England). 2016;143(24):4749–54.

37. Team RC. 2014.

38. Muzumdar MD, Tasic B, Miyamichi K, Li L, and Luo L. A global double-fluorescent Cre reporter mouse. Genesis (New York, NY : 2000). 2007;45(9):593–605.

39. Carroll TJ, Park JS, Hayashi S, Majumdar A, and McMahon AP. Wnt9b plays a central role in the regulation of mesenchymal to epithelial transitions underlying organogenesis of the mammalian urogenital system. Developmental cell. 2005;9(2):283–92.

40. Psychoyos A. Perméabilité capillaire et décidualisation uterine. C R Acad Sci. 1961;252:1515–7.

41. Lufkin H, Flores D, Raider Z, Madhavan M, Dawson M, Coronel A, et al. Pre-implantation mouse embryo movement under hormonally altered conditions. Molecular human reproduction. 2023;29(2).

42. Madhavan MK, DeMayo FJ, Lydon JP, Joshi NR, Fazleabas AT, and Arora R. Aberrant uterine folding in mice disrupts implantation chamber formation and alignment of embryo-uterine axes. *Development (Cambridge*, England). 2022;149(11).

43. Ma WG, Song H, Das SK, Paria BC, and Dey SK. Estrogen is a critical determinant that specifies the duration of the window of uterine receptivity for implantation. Proceedings of the National Academy of Sciences of the United States of America. 2003;100(5):2963–8.

44. Branham WS, Sheehan DM, Zehr DR, Ridlon E, and Nelson CJ. The postnatal ontogeny of rat uterine glands and age-related effects of 17 beta-estradiol. Endocrinology. 1985;117(5):2229–37.

45. Yamaguchi M, Yoshihara K, Suda K, Nakaoka H, Yachida N, Ueda H, et al. Three-dimensional understanding of the morphological complexity of the human uterine endometrium. iScience. 2021;24(4):102258.

46. Kurata R, Futaki S, Nakano I, Fujita F, Tanemura A, Murota H, et al. Three-dimensional cell shapes and arrangements in human sweat glands as revealed by whole-mount immunostaining. PloS one. 2017;12(6):e0178709.

47. de Paula F, Teshima THN, Hsieh R, Souza MM, Nico MMS, and Lourenco SV. Overview of Human Salivary Glands: Highlights of Morphology and Developing Processes. Anatomical record (Hoboken, NJ : 2007). 2017;300(7):1180–8.

48. Stewart CA, Stewart MD, Wang Y, Mullen RD, Kircher BK, Liang R, et al. Chronic Estrus Disrupts Uterine Gland Development and Homeostasis. Endocrinology. 2022;163(3).

49. Matsuo M, Yuan J, Kim YS, Dewar A, Fujita H, Dey SK, et al. Targeted depletion of uterine glandular Foxa2 induces embryonic diapause in mice. eLife. 2022;11.

50. Hancock JM, Li Y, Martin TE, Andersen CL, and Ye X. Upregulation of FOXA2 in uterine luminal epithelium and vaginal basal epithelium of epiERα-/- (Esr1fl/flWnt7aCre/+) mice†. Biology of reproduction. 2023;108(3):359–62.

51. Goad J, Ko YA, Kumar M, Syed SM, and Tanwar PS. Differential Wnt signaling activity limits epithelial gland development to the anti-mesometrial side of the mouse uterus. Developmental biology. 2017;423(2):138–51.

52. Hewitt SC, Li L, Grimm SA, Chen Y, Liu L, Li Y, et al. Research resource: whole-genome estrogen receptor α binding in mouse uterine tissue revealed by ChIP-seq. *Molecular endocrinology (Baltimore*, Md). 2012;26(5):887–98.

53. Ding T, Song H, Wang X, Khatua A, and Paria BC. Leukemia inhibitory factor ligand-receptor signaling is important for uterine receptivity and implantation in golden hamsters (Mesocricetus auratus). *Reproduction (Cambridge*, England). 2008;135(1):41–53.

54. Kholkute SD, Katkam RR, Nandedkar TD, and Puri CP. Leukaemia inhibitory factor in the endometrium of the common marmoset Callithrix jacchus: localization, expression and hormonal regulation. Molecular human reproduction. 2000;6(4):337–43.

55. Yang ZM, Chen DB, Le SP, and Harper MJ. Differential hormonal regulation of leukemia inhibitory factor (LIF) in rabbit and mouse uterus. Molecular reproduction and development. 1996;43(4):470–6.

56. Dhakal P, Fitzgerald HC, Kelleher AM, Liu H, and Spencer TE. Uterine glands impact embryo survival and stromal cell decidualization in mice. FASEB journal : official publication of the Federation of American Societies for Experimental Biology. 2021;35(10):e21938.

